# DeepC: Predicting chromatin interactions using megabase scaled deep neural networks and transfer learning

**DOI:** 10.1101/724005

**Authors:** Ron Schwessinger, Matthew Gosden, Damien Downes, Richard Brown, Jelena Telenius, Yee Whye Teh, Gerton Lunter, Jim R. Hughes

## Abstract

Understanding 3D genome structure requires high throughput, genome-wide approaches. However, assays for all vs. all chromatin interaction mapping are expensive and time consuming, which severely restricts their usage for large-scale mutagenesis screens or for mapping the impact of sequence variants. Computational models sophisticated enough to grasp the determinants of chromatin folding provide a unique window into the functional determinants of 3D genome structure as well as the effects of genome variation.

A chromatin interaction predictor should work at the base pair level but also incorporate large-scale genomic context to simultaneously capture the large scale and intricate structures of chromatin architecture. Similarly, to be a flexible and generalisable approach it should also be applicable to data it has not been explicitly trained on. To develop a model with these properties, we designed a deep neuronal network (deepC) that utilizes transfer learning to accurately predict chromatin interactions from DNA sequence at megabase scale. The model generalizes well to unseen chromosomes and works across cell types, Hi-C data resolutions and a range of sequencing depths. DeepC integrates DNA sequence context on an unprecedented scale, bridging the different levels of resolution from base pairs to TADs. We demonstrate how this model allows us to investigate sequence determinants of chromatin folding at genome-wide scale and to predict the importance of regulatory elements and the impact of sequence variations.

## Introduction

Mammalian genomes regulate gene expression through an intricate network of *cis* regulatory elements, such as enhancers, that can act over more then a million base pairs of DNA sequence. Enhancers have to physically interact with their target promoters to facilitate gene expression^1,2^. The accessible landscape of every regulatory element is in turn dictated by the three-dimensional genome organization, a hierarchical structure of insulated domains which vary between cell types and stages of development^3,4^. Genomic variants that alter chromatin architecture can lead to drastic misexpression^5,6^. Understanding chromatin architecture thus provides the basic scaffold for investigating the molecular mechanisms and predicting the functional implications of sequence and structural variants in health and disease.

Dissecting the factors underlying the formation and maintenance of genome structure requires high throughput, genome-wide approaches. However, assays for mapping chromatin interactions genome-wide such as Hi-C^7^, are expensive and time consuming. Extremely deep sequencing is required to map the chromatin architecture of a single cell type at high resolution. This severely restricts the usage for large-scale mutagenesis screens to identify the functional elements that underlie genome folding or for mapping the impacts of sequence or structural variants.

Computational models sophisticated enough to grasp the genomic determinants of chromatin folding would allow us to perform large-scale experiments *in silico.* The ideal chromatin interaction predictor should: 1) be sequence based, allowing us to map the determinants of chromatin folding and predict the impact of variation down to the base pair level; 2) incorporate large scale chromatin context, as window to window based approaches will miss key contextual information such as interjacent insulators; 3) be high resolution, to capture the intricate structures from insulated neighborhoods to megabase scale topologically associated domains (TADs); and 4) generalize well to unseen data, confirming the learned predictive features and improving confidence in variant effect predictions.

Although promising methods have been published, none addresses all these aspects simultaneously in a single framework. Methods that use information at the base pair level do not capture large scale continuous context and rather focus on window to window based predictions^8–10^. Methods that integrate a large genomic context do so by coarse segregation into genomic features^11–13^ or polymer beads^14–16^, thus compromising on the base resolution.

Here we present deepC, a deep neuronal network approach that can accurately predict Hi-C chromatin interactions from DNA sequence at megabase scale. DeepC uses Hi-C data to train a sequence based model that generalizes well to unseen data such as hold out chromosomes not used in training or validation. DeepC predicts chromatin folding at large scale and high resolution, bridging the gap from base pairs to TADs. We show that deepC works across cell types, learning cell type specific chromatin interactions. DeepC is applicable to a variety of Hi-C sequencing depths and can be used to fine map chromatin interactions even from low resolution Hi-C.

We validate deepC predictions with high resolution NG Capture-C^17^ at over one hundred loci. We then demonstrate how deepC allows us to investigate sequence determinants of chromatin folding at genome-wide scale and base pair resolution. We employ deepC to perform a genome-wide *in silico* screen, estimating changes in chromatin interactions upon deletion of every potential regulatory element. This allows us to dissect the average importance of promoters, enhancers and boundary elements and show the importance of all three of these elements in accurately predicting genome folding. Additionally, we demonstrate deepC’s utility for interpreting the effect of sequence and structural variants on chromatin architecture.

## Results

### A Deep Neural Network accurately predicts chromatin structure from DNA sequence

A model for accurate predictions of chromatin interactions from DNA sequence needs to meet two challenges dictated by the nature of chromatin folding. The model must include large-scale context but do so at base pair resolution to simultaneously capture large-scale structures such as TADs and smaller regulatory elements on the scale of 10s to 100s of bps and their intricate interplay.

To integrate these aspects, we built on a convolutional neuronal network architectures that have proven powerful for predicting chromatin features from ∼ 1 kb DNA sequence^18,19^ (Figure 1A). We combined this with a dilated convolutional network that excels at integrating large-scale context^20,21^. Our final model, deepC, operates on megabase scale DNA input while maintaining the input resolution at the single base pair.

**Figure 1.**
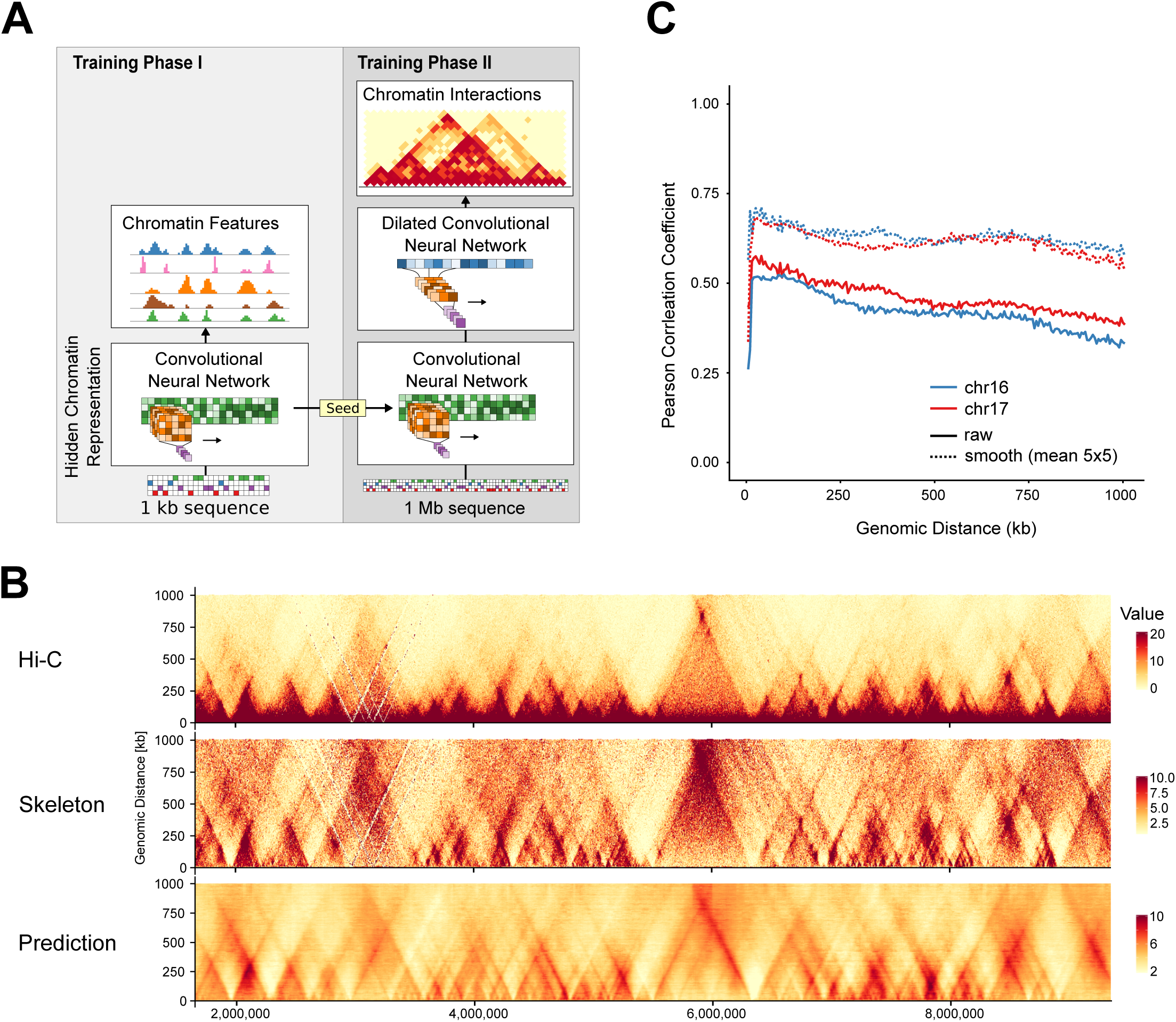
Predicting Hi-C interactions from DNA sequence. **A)** Overview of the deepC architecture and training workflow. **B)** Overlay of Hi-C data, the derived Hi-C skeleton and the interactions predicted from DNA sequence using deepC. Shown is a ∼ 7 Mb region on one of the test chromosome 17. **C)** Distance stratified Pearson correlation between the Hi-C skeleton and the deepC predictions (solid lines). Shown are both test chromosomes (16, 17). Because the Hi-C skeleton is noisy, a small 5×5 mean filter was applied to the skeleton and the correlation with the deepC predictions measured (dotted lines).

When optimizing the network architecture and data encoding (Figure 1B, Supplementary Figure 1A), we found two factors crucial for successful learning and generalization: First, we normalize the proximity signal in Hi-C data by percentile normalizing the interactions stratified over different genomic distances (Supplementary Figure 1B, for details see methods). Effectively this skeleton encoding reveals the informative longer range interactions underlying Hi-C data. Second, we found a two-step training process to yield much better results (Fig 1A, Supplementary Figure 2). We first train a convolutional neural network to predict a compendium of chromatin features (eg, open chromatin regions, CTCF binding site etc.). We then use the hidden layers of the trained network to seed the first layers of our chromatin interaction network. The full network then simultaneously optimizes the initial layers and learns to combine them through the dilated convolutional layers to predict chromatin interactions. This can be conceptualized as transfer learning, where the network learns to recognize chromatin features from DNA sequence first and then learns to use the learned underlying sequence patterns and combine them over large distances for chromatin interaction prediction. We observed fast convergence when seeding with a pre-trained chromatin feature network.

DeepC achieves accurate predictions over hold out chromosomes (Figure 1C). The average distance stratified Pearson correlation between the Hi-C skeleton and deepC prediction is ∼ 0.44 and ∼ 0.62 when applying a small mean filter on the noisy skeleton data (Figure 1D). This is comparable to orthogonal methods that predict chromatin interactions directly from cell type specific chromatin features^11–13^, although genome folding prediction from sequence is a much more difficult task.

Visually, deepC yields smooth but intricate predictions that resolve the hierarchical nature of TADs and insulated domains at high resolution. This resolution is only matched by Hi-C data with very deep sequencing depth (GM12878 primary ∼ 3.6 B reads). When we apply deepC to less deeply sequenced Hi-C data (e.g. K562 ∼ 1.3 B reads), the predicted resolution surpasses that of the used Hi-C data itself, effectively fine mapping Hi-C interactions (Supplementary Figure 3).

We applied deepC over seven cell types with different sequencing depths^22^ (Supplementary Figure 4) and across Hi-C binning sizes (Supplementary Figure 5). Using sub-sampling experiments we show that deepC works robustly over less deeply sequenced Hi-C and may be used to boost their resolution (Supplementary Figures 6, 7). Importantly, deepC is able to learn tissue specific chromatin interactions (Supplementary Figure 8). Furthermore, the combination of skeleton transformation and the smoothness of deepC predictions allows us to call domain boundaries at higher resolution then possible from raw Hi-C data using an insulation score based approach (Supplementary Figure 9, see Methods).

### Validation of high-resolution domain structure

To validate the deepC predicted Hi-C structures and derived boundaries, we performed NG Capture-C from 220 viewpoints in two cell types (GM12878 and K562). NG Capture-C probes the chromatin interactions from viewpoints of interest at high resolution and sensitivity, allowing us to validate deepC’s fine grained domain predictions. We specifically target loci where deepC predictions show more detail then the corresponding Hi-C data. Half the viewpoints were chosen from validation and test chromosomes that have been held out from training. We targeted 81 CTCF sites to see specific looping events and 139 sites within insulated domains but avoiding active elements, to detect domain boundaries.

NG Capture-C does resolve the interaction domains and specific interactions within at greater detail then the Hi-C skeleton or deepC predictions (Supplementary Figure 10). However, we observe good agreement between NG Capture-C tracks and the predicted Hi-C structure (Figure 2A). The domain boundaries predicted from deepC overlap with spikes in the CTCF capture, corresponding to CTCF loops and spikes and drops in the intra domain capture, highlighting the interaction domain boundaries. To quantify this, we called domain boundaries from the deepC predictions at high resolution and we observe a clear enrichment of distance normalized NG Capture-C signal over the predicted boundaries (Figure 2B). This enrichment was seen in both types of viewpoints and was strongest in the CTCF captures due to the strength of CTCF to CTCF loops.

**Figure 2.**
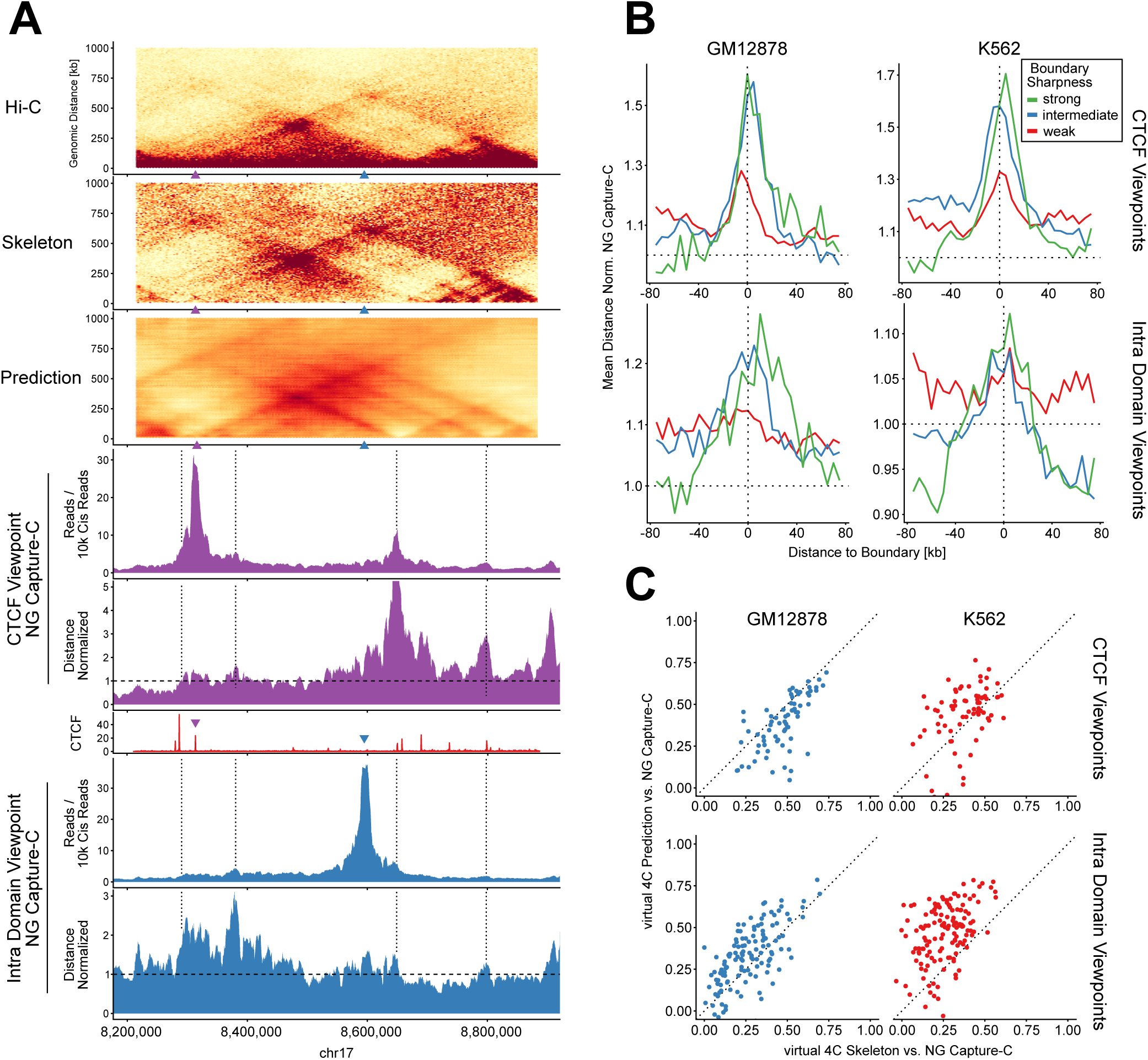
Capture-C validation of deepC predictions from 220 viewpoints. **A)** Example region with overlay of GM12878: Hi-C, skeleton and deepC prediction; NG Capture-C tracks, distance normalized NG Capture-C tracks and CTCF ChIP-seq track (red). Shown is a CTCF viewpoint (purple triangle) and an intra domain viewpoint (blue triangle) not overlapping with any active elements. Dotted lines are aligned to predicted boundaries for orientation. Dashed lines in distance normalized Capture-C tracks indicate the expected interaction value. **B)** Meta profile of the average NG Capture-C signal over domain boundaries called from the deepC predicted interaction map at high resolution. Boundaries were classified based on their sharpness (see Methods). Only NG Capture-C interactions from viewpoints within 1 Mb of the respective boundary were used. **C)** Comparing the correlation between the validation NG Capture-C profiles and the virtual 4C profiles derived from the Hi-C skeleton (x-axis); and the deepC prediction map (y-axis) from all viewpoints in two cell types.

Using virtual4C, we can predict the interaction profiles of our capture viewpoints from Hi-C data and deepC predictions (Supplementary Figure 10). When we correlate the NG Capture-C profiles with the virtual4C maps derived from Hi-C skeleton and the deepC predictions respectively, we find good correlation (Figure 2C). This was robust across training, test and validation chromosomes and across A and B compartments (Supplementary Figure 11). However, virtual4C does not predict the high resolution NG Capture-C profile perfectly. This is true regardless if the virtual4C was derived from the Hi-C skeleton or the deepC predictions. Remarkably, both methods are competitive although deepC makes these predictions from sequence rather then sampling from actual chromatin interaction data requiring billions of reads.

### Predicting the impact of variation on chromatin structure

Genomic variants, such as structural variations can alter gene expression by rewiring regulatory interactions such as enhancer – promoter contacts^1,23,24^. Such variants and their associated links to gene dysregulation have been shown to be drivers of severe phenotypes and disease^6,25,26^.

DeepC allows us to predict the impact of sequence variation. DeepC’s base pair resolution and large-scale context allow us to predict the impact of single nucleotide variants and indels as well as larger structural variants in the same framework. In Figure 3A we demonstrate the predicted impact of deleting a single CTCF site. DeepC predicts the merging of subTADs and formation of new subTAD boundaries at different CTCF sites. Calculating the difference between the predicted chromatin structures we can summarize the predicted impact of a mutation into a single score.

**Figure 3.**
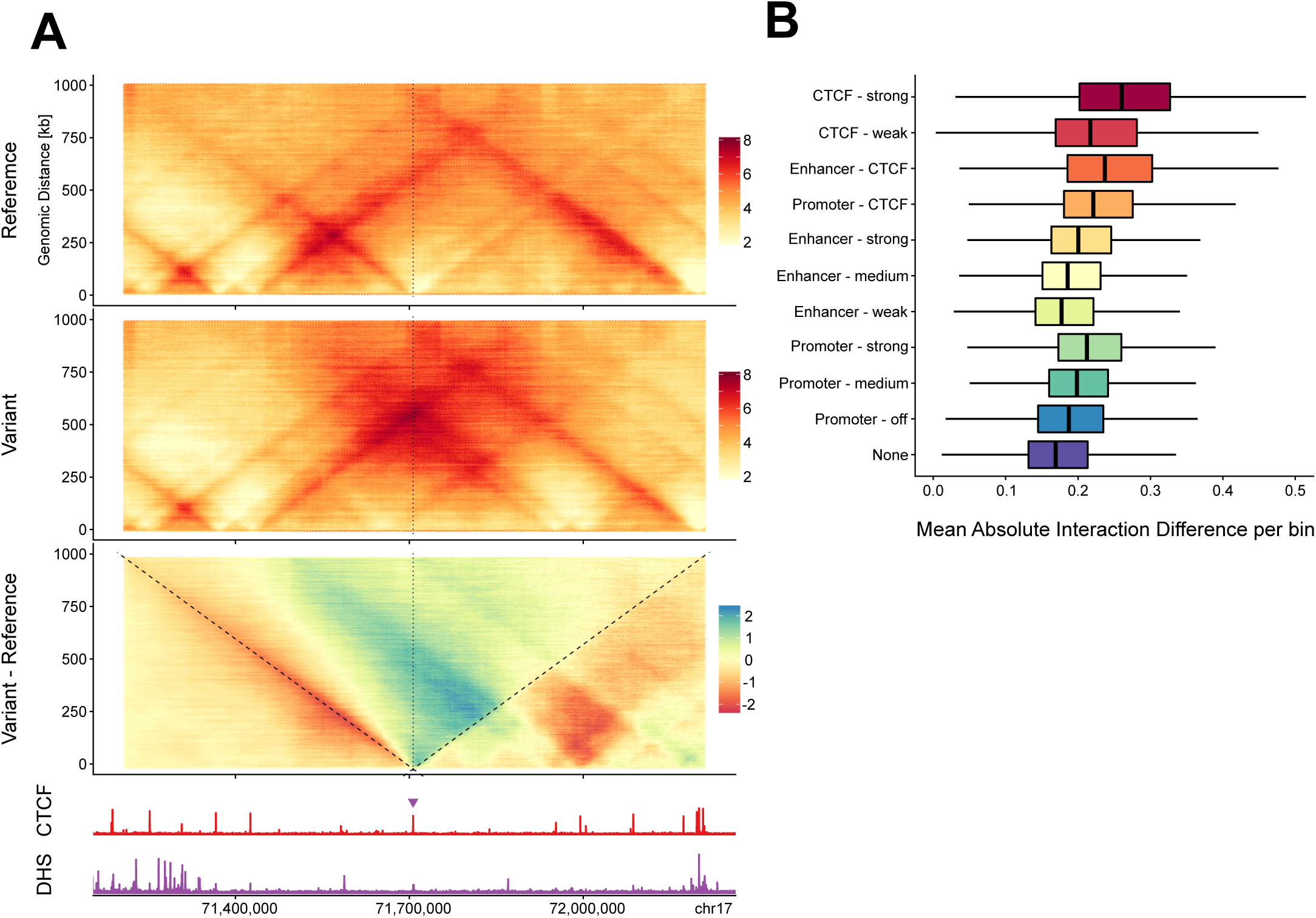
Genome wide in silico deletion screen of open chromatin and CTCF binding sites. Using the GM12878 model, all open chromatin and CTCF binding sites were individually deleted in silico and the predicted impact on chromatin structure quantified. **A)** Example prediction of the impact of deleting a single CTCF site (purple triangle) on chromatin structure. Shown is the reference deepC prediction, the predicted interaction upon deletion and the difference between both maps. GM12878 CTCF ChIP-seq and DNase-seq tracks are aligned underneath. The deepC prediction values were bounded between 2 and 8 for contrast. **B)** Summary of the predicted chromatin interaction differences between reference and variant. Differences were quantified as the mean absolute interaction difference per bin to bin interaction. Each deletion was classified based on the overlap with genome segmentation classes. Outliers not plotted.

A long standing question has been which functional elements within the genome underlie the patterns of genome folding seen in data such as Hi-C and NG Capture-C. To investigate this we used deepC to perform a serial *in silico* deletion of all active elements genome wide to determine their importance to chromatin interactions (Figure 3B). As expected, we find CTCF sites as well as enhancers and promoters with proximal CTCF binding to have the strongest average impact. Interestingly, we find promoter deletions to have on average a stronger impact then enhancer deletions. Within promoters and enhancers respectively, we find the ones strongly marked by activity associated histone marks to have a stronger predicted impact. Our observations match with insights from orthogonal methods, that found CTCF binding and cell type specific active chromatin marks^12,13^, and CTCF binding and RNA expression^11^, to be most predictive of chromatin interactions, respectively. Considering the fidelity of the deepC predictions this suggests that the patterns of 3C genome folding seen in methods such as Hi-C arise from the interplay of the activities of enhancers and promoters as well as CTCF bound sites.

### Probing sequence determinants of chromatin folding at base pair resolution

The base pair resolution of our model allows us to interrogate the sequence determinants of chromatin folding up to megabase scale at the base pair level. Using deepC we can calculate the relative importance of every base pair for predicting chromatin interactions up to a megabase away. This importance score, called saliency, quantifies how much the chromatin interaction prediction would change, if we change a single base pair. Explicitly, the saliency score measures the gradient of the chromatin interaction predictions with respect to the sequence input.

The IKZF2 locus (Figure 4A, B) exemplifies the saliency patterns we observe genome-wide (Figure 4B). The saliency peaks sharply at CTCF sites, but we also find broad saliency peaks frequently located underneath promoter regions. In comparison, enhancers only show strong saliency enrichment when they are co-localized with CTCF sites (Figure 4C). Since a high saliency score can tag existing determinants for chromatin folding as well as potentially influential regions upon mutation, we also observe saliency peaks under unbound potential CTCF motifs. When we visualize the saliency score at base pair resolution we recover CTCF and other transcription factors binding motifs. Interestingly, when we compare the saliency scores derived from the GM12878 and the K562 model (Figure 4B, D), at genes with differential expression, namely IKZF2 strongly expressed in EBV transformed lymphocytes and TAL1 strongly expressed in erythroid cells, their promoter regions show differential saliency enrichment.

**Figure 4.**
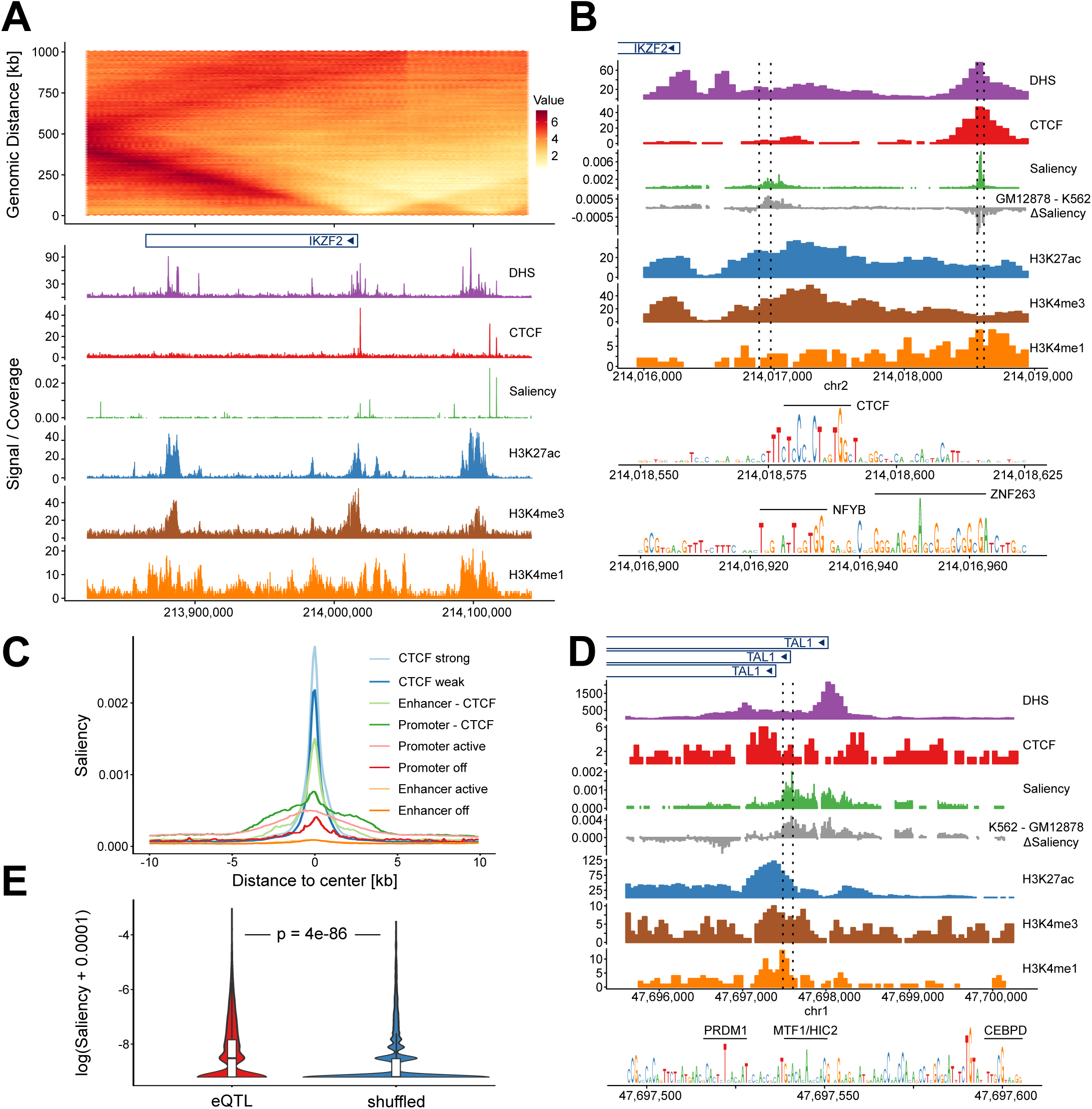
Mapping sequence determinants of chromatin folding at bp resolution. Overlay of deepC chromatin interaction prediction for GM12878 with matching chromatin data: DNase-seq and ChIP-seq for CTCF and chromatin marks. At gene scale **(A)** and promoter scale **(B)**. Shown in green is the saliency score which is a proxy for the importance every base has in predicting the chromatin interactions of that region. The saliency score shows sharp peaks overlapping CTCF binding sites and broader peaks overlapping active gene promoters as seen on the meta plot of average saliency per chromatin element class **(C)**. Resolving the saliency score at base pair resolution **(B)** highlights CTCF motifs and general transcription factor binding motifs (regions within dotted lines, CTCF site right and top, promoter left and bottom). When subtracting the saliency scores derived from the K562 deepC model from the GM12878 model (grey) the IKZF2 promoter shows enriched saliency signal in GM12878. The reverse is true for the TAL1 promoter in the K562 model **(D)**. This matches the tissue specific expression patterns of IKZF2 and TAL1. Note that CTCF sites show a difference in the tissue specific saliency score because both models have been trained independently. **E)** GTEx derived eQTLs specific for GM12878 cells that intersect with GM12878 open chromatin sites are enriched for higher deepC saliency scores compared to an equally sized, reshuffled set of SNPs within open chromatin. This difference is significant (one sided Kolmogorov–Smirnov Test). It suggests that eQTLs are enriched for SNPs at positions that influence 3D structure.

As bases with high saliency scores mark important positions for chromatin architecture, we hypothesized that they are likely to affect gene expression. To test this, we retrieved GM12878 cell type specific eQTLs (GTEx v7). We selected the ∼5000 eQTLs that are located in open chromatin (DNase-seq) and or CTCF sites (CTCF ChIP-seq, ENCODE) and are thus likely to lie in regulatory elements. These eQTLs have significantly higher saliency scores than SNPs randomly re-sampled from open chromatin and CTCF sites (Figure 4E). Practically, the saliency score fine maps accessible elements with respect to impact on chromatin architecture.

## Discussion

Mammalian chromatin architecture unfolds at megabase scale constraining distal regulatory interactions within TADs and smaller insulated domains^4,5,27^. Ultimately, chromatin interactions are encoded in the DNA sequence through an intricate interplay of protein binding sites and other sequence determinants. Bridging the gap from the single base pair level to large-scale chromatin interactions is a key challenge for understanding chromatin architecture and it’s interplay with gene regulation^28,29^. We developed deepC as our approach to traverse this gap. DeepC is the first sequence based deep learning model that predicts chromatin interactions from DNA sequence within the context of the megabase scale of the genome. We believe this large scale context integration is crucial for predicting chromatin interactions, to determine their sequence determinants and for interpreting the impact of sequence variations.

Other sequence-based methods^8,9^ only predict interactions between two DNA windows. This approach neglects the influence of surrounding and interspersed elements. For example, the presence of a strong (TAD) boundary in between windows that dictates the chromatin interaction landscape of the locus will be missed. Recent methods also predict chromatin interactions from cell type specific chromatin features, such as CTCF and histone modification ChIP-seq, DNase-seq and RNA-seq. These methods capture the chromatin features between two interaction windows by: averaging them^10,12^; or by capturing the number, distance and orientation of chromatin features such as CTCF through careful feature engineering^11^. However, these methods are not sequenced based, limiting their ability to map the determinants of chromatin folding at high resolution. To predict the impact of smaller sequence variants these models must also map the variant impact on chromatin features through experiments or preceding predictive models. In contrast, deepC offers an end-to-end training and prediction approach. Polymer models have also shown a strong potential for predicting the impact of structural variations on chromatin folding^16^. However, the underlying polymer model is optimized for every chromosome, based on the corresponding Hi-C data rather than from sequence and so predicts the impact of a variation on the same region used to train the polymer model. Thus, polymer models try to answer a different question, because they do not attempt to generalize to unseen regions. Furthermore, polymer models learn to encode chromatin as string of beads representing DNA windows, potentially down to the resolution of individual factor binding sites but not DNA sequence. Predicting the impact of single nucleotide variants and small indels would thus require intermediate models.

Our current deepC models learn to predict chromatin interactions of a single cell type at a time. Therefore comparisons between models, such as between cell types, need to distinguish between stochastic variability in the training process and biologically meaningful cell type specific features. To facilitate the discovery and interpretation of tissue specific factors of chromatin folding, we work towards a single generalized network that encodes cell type identity and learns to predict chromatin interactions across tissues in a single process. Although, using megabase DNA context encoded at base pair resolution is unprecedented, deepC does not yet capture interactions greater than the megabase scale. However, this limitation purely reflects current computational limits and inevitable improvements in hardware and further optimizing our model architecture will allow deepC to use larger sequence context in the same framework. In addition, we found deepC models to yield substantially better predictions, when being pre-seeded with hidden layers optimized to predict a compendium of chromatin features in a first training phase. Although these layers are refined in the second, chromatin interaction, training phase, the network remains biased towards recognizing the sequence determinants of chromatin features included in the first phase. Predictive sequences for chromatin features not included or potential sequence determinants that directly influence chromatin folding may still be learned by reusing and refining existing hidden layers, but are learned in a less efficient manner. We believe that future improvements in model architecture as well as data availability will allow us to address these limitations in a more efficient manner.

We believe deepC will prove to be valuable tool for dissecting the 3D regulatory genome code. We provide trained deepC models for seven cell types. Users can utilize these models to help fine map chromatin interactions, interrogate the features and sequence determinants of chromatin folding and predict the impact of variation *in silico*. Users can further train deepC models on new Hi-C data to derive novel cell type models.

We demonstrated some of the diverse use cases for employing deepC. Our genome wide deletion screen of potential regulatory elements confirmed that CTCF site deletions to be most likely to cause strong chromatin interaction changes. Furthermore, deepC indicates both promoters and enhancers to contribute to genome folding even in the absence of CTCF. Interestingly, we identified promoter deletions to have a higher predicted effect then enhancer deletions. Comparing enhancers and promoters respectively, we find the deletions of elements associated with active chromatin marks to have higher predicted impacts. Our observations are in line with findings from orthologous methods, that find CTCF binding, open chromatin, active histone marks and RNA-seq to be most predictive^10–13,16^. Interestingly deepC consistently predicts domain like structures with frequent and pronounced stripes rather then distinct point wise interactions. Moreover, deepC highlights active regulatory elements to be important for predicting chromatin interactions, even when they are not co-localized by CTCF binding sites. These observations support a loop extrusion mediated model of chromatin folding ^15^ over a model of pair wise chromatin interactions. Furthermore, we fine mapped the sequence determinants of chromatin architecture at base pair resolution and linked this information with effects on gene expression.

We believe deepC presents a valuable tool for interrogating chromatin architecture that will facilitate future research. Furthermore, deepC represents an important step towards predictive models of gene regulation that integrate the intricate and long ranged chromatin landscape of mammalian genomes.

## Methods

### Chromatin feature data

As chromatin feature compendium the ENCODE^30^ and Roadmap^31^ chromatin data utilized in DeepSEA^18^ were used. Narrow peak calls (hg19) for 918 experiments were downloaded. The data was supplemented with additional erythroid linage data. Five sets of ATAC-seq data from Corces et al.^32^, two DNase-seq experiments published earlier^33^ and ten ATAC-seq and one CTCF ChIP-seq experiment from in house erythroid differentiations were used. All data used are listed in Supplementary Table 1. All additional data, was aligned to hg19 using the NGseqBasic pipeline^34^. Peaks were called with macs2 (default parameters, -q 0.01). The peak signals were aggregated following the procedure described in Zhou et al.^18^. In brief, the genome was split into 200 bp bins. All peak calls were intersected with these bins. If a bin overlaps a peak call to at least 50 % (100bp), the bin is labeled as belonging to that dataset class. A peak can have as many classes as peak calls used. All genomic bins that do not intersect with at least one peak call were discarded. Only autosomes were used for all analysis.

### Hi-C data

The publicly available, deeply sequenced Hi-C data from Rao et al.^22^ was used. The available 5 and 10 kb resolution intra chromosomal contacts maps of 8 cell lines were downloaded and normalized using the provided KRnorm factors. In addition, the primary GM12878 replicate was realigned from raw fastqs using HiCPro ^35^. HiC-Pro was run with default parameters but modified to allow for minor multi-mapping in duplicate regions such as globin genes. Bowtie2 was run with the -k option set to 3 and filtering reads with more then 2 multimapping positions. Effectively this mimics a bowtie (v1) run with flag -M 2 set. The passing reads were filtered for valid interactions and contact matrices built and ICE normalized using the HiC-Pro default. For the sub sampling analysis, the valid interactions from the realigned, primary GM12878 set were sub sampled to achieve 1 billion, 100 million and 10 million valid interactions, respectively.

### Hi-C encoding for deep learning

The genome was divided into bins matching the bin size of the respective Hi-C data resolution used for training (1 kb, 5 kb, 10kb). For every stretch of DNA of size 1 Mb + bin size bps (e.g. 1005000 for 5kb bins), the chromatin interactions associated with the window were assigned as squares in a vertical zig-zag pole over the center of the sequence window (see Supplementary Figure 1A). Every square encodes the Hi-C interactions observed between two bin sized windows of increasing distance (up to 1 Mb away). By sliding the large DNA stretch over a chromosome with a bin sized increment, this encoding recovers the chromosome wide Hi-C map up to an interaction distance of 1 Mb.

The Hi-C interaction signal was normalized to remove the linear distance dependence. The resulting signal can be regarded as the skeleton underlying the Hi-C signal. The Hi-C signal is normalized using a distance stratified, percentile normalization adapted for the unique nature of Hi-C data. The Hi-C data is percentile normalized across a chromosome separately for every interaction distance in bin sizes. It is of particularly interested to resolve high levels of Hi-C interactions at high resolution and only a low percentage of chromatin interactions is expected to yield strong interactions at larger distances for example the corners of TAD triangular structures. The percentile normalization was designed better resolve these high interaction levels at larger distances by using uneven percentiles in a pyramid like scheme (from low to high: 2× 20 %, 4 × 10 %, 4× 5 %, see Supplementary Figure 1B). The identifier of the respective pyramid percentile (1 – 10) was stored. The chromatin interaction network was then trained to predict the pyramid percentile identifier as a regression problem (see below).

### Deep neural network architectures

For predicting chromatin interactions with deep learning, we found a two step training process with transfer learning (Supplementary Figure 2) to yield substantially better results. First a convolutional neural network was trained to predict chromatin features from 1 kb of DNA sequence, using the compendium of 936 datasets described above. A general introduction into the use of convolutional neural networks for genomics applications can be found in^36,37^. The principle network architecture was adapted from DeepSEA^18^. Five convolutional layers were used (instead of 3 as in DeepSEA). ReLU was used activation function, followed by 1D max pooling and dropout at every convolutional layer. The five pooling layers were designed to pool a total of 1 kb into a single entry along the sequence axis. This greatly reduces the number of parameters to learn in the final, fully connected layer. Sigmoid activation was used for the fully connected layer to output an individual probability for each class (multi-label classification). The network parameters were trained by minimizing the sum of the binary cross entropies using the ADAM optimizer in batches of 100. Hyperparameters were optimized by grid search. For details on the final, best performing hyperparameters see Supplementary Note. We did not find batch normalization to benefit convergence speed or robustness in our setting.

In the second step a chromatin interaction network was used to predict Hi-C interaction from DNA sequence. The chromatin interaction network takes as input 1 Mb + 1x Hi-C bin size [bp], (e.g. 1005000 for a 5 kb bin network). The first module consists of five convolutional layers, with ReLU, max pooling and dropout. Dimensions and hyperparameters for this stage were equal to the chromatin feature network. The hidden weights were initialized by seeding with the weights of the trained chromatin feature network from step one.

The second module is a series of nine dilated, gated convolutional layers with residuals (see Supplementary Notes). Following PixelCNN^38^ and WaveNet^20^, gated convolutional layers require training double the amount of filter parameters, but have the potential of modeling more complex functions through their multiplicative units, similar to LSTMs. The residual units allow information to propagate more easily through the network without having to necessarily pass through convolutions^39^. The dilation rates were chosen to reach the full sequence context in the last layer. GPU memory limited us to using a batch size of 1. The dilated layers were followed by a fully connected layer. The output of the fully connected layer are the predicted interaction strengths (in units matching the pyramid percentile normalization). The model was trained with ADAM to minimize the mean square error between the outputs and the true percentiles. Hyperparameters not dictated by the seeding procedure were optimized using grid search.

### Training the deep neural networks

For both training procedures the data were split into training, validation and test set based on chromosomes. For the chromatin feature network chr11 and 12 were used for for validation and chr15, 16 and 17 for testing. For the chromatin interaction network, to increase the number of training examples the same validation chromosomes were used but only chr16 and 17 were used as test chromosomes. All models were trained on NVIDIA Titan V cards. Fully training the chromatin feature network required 14 epochs. The training set order was reshuffled after every epoch.

To minimize the amount of times large chunks of DNA sequence have to be loaded into memory the network is trained on one chromosome at a time. Within a chromosome, the order in which training batches are drawn is random. Interestingly, we observed that the chromatin interaction network, when seeded with the pre-trained weights in the first convolutional filters, converges quickly, after training on ∼ 3 - 6 chromosomes and only marginally improves after training for an entire epoch or longer. Models were trained for one full epoch as we have not observed significant improvement after training for longer and the limited batch size as well as the network complexity make training slow.

### Predicting chromatin interactions and estimating interaction changes

Using a trained model the chromatin interactions of a region of interest or of the entire hold out test chromosomes can be predicted from sequence. By modifying the input sequence the chromatin interaction changes caused by a sequence variants can be predicted. For calculating differences in chromatin interactions, the chromatin interactions over the reference sequence was predicted for every position that is within 1 Mb (plus 1x Hi-C bin size) of the sequence variant. This matches the respective models spatial reach. The reference sequence is then modified to match the sequence variant of interest. After predicting the chromatin interactions over the variant sequence, the difference is quantified by calculating the absolute difference between reference and variant prediction. Then the mean of the absolute difference over all covered interaction tiles (bin sized interaction windows) can be calculated. This is straight forward for single or multiple base pair substitutions as long as the same number bases between reference and variant match up. For variants like deletions, there are to options for modeling the variant. 1) The reference sequence can be deleted. This requires to add a matching number of additional base pairs (e.g. at the downstream end) to maintain the total sequence length for the model. This has the benefit of modeling the actual variant sequence and captures effects at the breakpoints, e.g. creating a new binding site. The drawback is that all chromatin interaction predictions are downstream of the variant are “moved” towards the center. For larger deletions (∼ >100 bp), this has the effect of creating a shadow of chromatin interaction changes. In variants that have a strong effect on chromatin structure, the shadow effect is small compared to the distinguishable variant effect that we are actually interested in. In contrast, in variants that don’t seem to have a predicted impact, the shadow effect is the only observable feature. 2) Alternatively, the affected bases can be changed to N’s instead of deleting them. This removes the shadow effect but may loose breakpoint effects. The same principle translates to insertions. In practice, the only difference we observe between both approaches is for visualization with the shadow only being present when using method 1). See Supplementary Figure 12 for an example impact of choosing a deletion mode. For estimating the impact of a deletion and for the genome wide deletion screen below approach 1 was used. To remove the shadow effect, approach 2 was used for visualizing the impact of larger deletions.

### Distance Stratified Correlation

We can calculate the correlation between the Hi-C skeleton and the deepC predicted interaction on the hold out test chromosomes. For calculating the correlation between Hi-C maps, e.g. from replicates, without accounting for the distance dependence signal leads to inflated correlation values^40^. To mitigate this effect, it is possible to stratify the correlation over the distance between interacting regions^40,41^. The Hi-C skeleton transformation employed already accounts for the distance dependence in terms of DNA polymer proximity. Nevertheless, we still might observe differences between predictive performance at different linear distance. For example, a model might be better at predicting short ranged or intermediate contacts over long ranges ones. To monitor such behavior, the correlation between the model predictions and the Hi-C skeleton was stratified over linear distance in bins matching the encoded bin size. The Pearson correlation coefficient was calculated between the Hi-C skeleton percentile tag (1 – 10) and the predicted regression score. Note that the skeleton percentiles are discrete, while the regression score is continuous. The Hi-C skeleton is noisy even at very deep sequencing depths (e.g. GM12878). Therefore a small mean filter was employed using a 5×5 bin window to smooth the skeleton. The distance stratified correlation was calculated between the prediction and the raw or the smoothed skeleton respectively.

### Selection of Validation Capture probes

A total of 220 viewpoints was selected for validating the deepC predictions. We specifically selected genomic locations where the Hi-C data and deepC predictions differed in detail or where the structures deepC predicted where only very faintly noticeable in Hi-C data. Two sets were designed, one targeting CTCF sites and one set targeting intra domain viewpoints that lie within a distinct Hi-C/deepC domain but are not intersecting with any potential functional elements. In total, 81 CTCF sites and 139 intra domain viewpoints were selected. Capture probes were designed using CapSequm (http://apps.molbiol.ox.ac.uk/CaptureC/cgi-bin/CapSequm.cgi), filtering out repetitive probe regions as described in the online documentation. For the final probe design see Supplementary Table 2.

### Cells, cell culture and fixation

Human GM12878 lymphocyte cell line, were obtained from the NIGMS Human Genetic Cell Repository at the Coriell Institute for Medical Research and cultured in RPMI 1640 supplemented with 15% FBS, 2mM L-Glutamine and 100U/ml Pen-Strep at 37 °C in a 5% CO2 incubator. K562 cells were supplied by the WIMM transgenics facility. Cells were maintained in RPMI 1640 media supplemented with 10 % FCS at 37 °C in a 5% CO2 incubator. Both cell types were fixed and processed using the same protocol. Cells were resuspended at 1×10^6^ cells per ml and fixed at room temperature with 2% v/v formaldehyde for 10 minutes. Fixation was quenched with 120 mM glycine. Cells were washed with ice cold PBS. Cells resuspended in cold lysis buffer (10 mM Tris, 10 mM NaCl, 0.2% Igepal CA-630 and complete proteinase inhibitor (Roche)) and snap frozen to −80 °C.

### Capture-C

3C libraries were prepared as previously described^17^ with the following modifications: Centrifugation’s were performed at 500 rcf, thermomixer incubations were set to 500 rpm, and following ligation chromatin was pelleted by centrifugation (15 min, 4°C, 500 rcf) and the supernatant discarded. Double capture was performed as described previously^17^ with biotinylated oligonucleotides (IDT xGen Lockdown Probes) in two pools (Supplementary Table 2) with 3 pg of each oligonucleotide per 3C library. The generated Capture-C libraries were sequenced using Illumina sequencing platforms (V2 chemistry; 150-bp paired-end reads). To resolve even subtle changes in chromatin interaction domains at high resolution, the libraries were deeply sequenced (GM12878 CTCF - 128 M; GM12878 intra domain −118 M reads; K562 CTCF – 302 M; K562 intra domain – 289 M reads).

### Capture-C analysis

Capture-C data were mapped, quality controlled and visualizing using CCseqBasicS (https://github.com/Hughes-Genome-Group/CCseqBasicS) following the procedure described previously^17^.

### Distance normalize Capture-C tracks

To compare them to the Hi-C skeleton, Capture-C tracks were normalized for distance dependence per viewpoint. The number of interactions with each restriction enzyme fragment was extracted and the distance to the viewpoint recorded. The interactions were then normalized for the total number of cis interactions for the respective viewpoint. Pooling this information across all viewpoints we observed that the distance decay approximately follows a log – log linear trend when we split the data into three distance bins (close, intermediate and far). The distance thresholds for these bins were empirically optimized for every Capture-C set (see Supplementary Table 3), excluding everything closer then 2.5 kb to the respective viewpoint. The distance decay is then approximated by three linear regression fits, one for each distance bin. The distance normalized interactions were calculated per viewpoint by dividing the observed *cis* normalized interactions with the expected interactions from the linear fit at the respective distance.

### Insulation score boundary calling

Interaction domain boundaries were called on the deepC predicted interactions using an insulation score based approach that was adapted from the procedure described by Crane et al.^42^. Using the 5 kb bin sized data, the mean insulation score profile was calculated based on a 25 kb window to allow for a more intricate boundary call. The first derivative, or delta vector, of the insulation score profile was approximated using a 1D Sobel operator. Zero crossings in this delta vector represent local minima and maxima of the insulation score. Maxima were discarded. The remaining boundaries were further filtered by calculating the approximation of the second derivate of the insulation profile using the same procedure described above. The height of this delta2 vector reflects the change in insulation score change, with sharper boundaries having a higher delta2 score. Boundaries with a delta2 score smaller then 0.1 were removed and the remaining boundaries were stratified based on their delta2 score.

### Distance normalized Capture-C signal over boundaries

The mean normalized Capture-C signal over boundaries called from deepC interaction predictions was calculated. In Capture-C tracks from single viewpoints, domain boundaries can be subtle and get harder to detect the further away from the viewpoint they are located. We therefore focused on interactions in a moderate distance to the viewpoint and excluded all interactions that are more then 1 Mb away. The mean normalized Capture-C signal over predicted boundaries was visualized as profiles, that were stratified for boundary sharpness using the delta2 score described above.

### A/B compartment call

Hi-C A and B compartments were called using the cscore approach^43^. The cscore orientation corresponding to A and B compartments were assigned per chromosome based on the overlap with active chromatin marks.

### Virtual4C from Hi-C skeleton and deepC maps

By extracting all interacting windows with a viewpoint of interest, Hi-C data can be transformed into virtual 4C profiles. For this work, virtual4C profiles were derived from the Hi-C skeleton and the deepC predictions, thus yielding distance dependence normalized profiles. Virtual4C profiles from the Hi-C skeleton and deepC predictions were compared to distance normalized Capture-C tracks by calculating the respective Pearson correlation of all interactions within 1 Mb from a given viewpoint. Note that the skeleton percentiles are discrete, while the deepC score and Capture-C tracks are continuous. In comparison, virutal4C profiles from the skeleton are relatively sparse even at deep sequencing depth.

### Saliency score

The saliency was calculated as the gradient of the model output, the predicted chromatin interactions, with respect to the sequence input. The saliency score was calculated at bp resolution. At a single step the saliency score relates to the vertical interaction pole on the center of a given sequence window. The sequence window is then moved in bin sized steps and the saliency is averaged over all sequence windows (sized 1 Mb + bin size) that include the respective base pair. For easy of visualization and interpretation the absolute value of the saliency score were used. Saliency scores derived from the 5 kb resolution GM12878 and K562 model were used for visualization, respectively.

### Chromatin segmentation

GM12878 and K562 chromatin data were downloaded from the ENCODE data portal (see Supplementary Table 4). Filtered alignments to hg19 were downloaded and replicates were merged. Peaks were called using macs2 with default settings and -q 0.01. Deeptools was used to create bigwig coverage tracks. No normalization was used. DNase-seq and CTCF ChIP-seq peaks were merged to a union set merging peaks within 10 bp of each other (bedtools merge -d 10). Union peaks were formated to 600 bp elements centered on the peaks. Deeptools was then used to extract the read coverage for each chromatin dataset over each peak union element. For this elements were extend to 1000 bps to better capture flanking histone modifications. Using the derived count matrix, chromatin classes were segmented using GenoSTAN running on the elements rather then entire chromosome stretches. The HMM model was trained using the Poisson log-normal distributions. Twelve classes were fitted and merged into 11 classes based on similarity of the chromatin signatures. The classes were manually curated and classified into promoter, enhancer and CTCF sites with varying activity levels based on H3K27ac.

### eQTL data analysis

EBV transformed lymphocyte specific eQTLs were retrieved from GTEx (v7 accessed from the GTEx portal 01/03/2019). A union of DNase-seq and CTCF ChIP-seq peaks was created using bedtools merge. The eQTL SNPs were filtered for intersection with the union of GM12878 open chromatin and CTCF peaks. Indels were removed. A background SNP set was constructed by shuffling the eQTL SNPs on the respective same chromosome and forcing them to stem from within the union peaks (bedtools shuffle -chrom -incl). Absolute saliency scores of the SNP bases derived from the 5 kb resolution GM12878 model were extracted. Empirical cumulative distributions were derived and tested for significance using a two sample Kolmogorov-Smirnov test (R, ks.test, reshuffled SNP saliency vs. eQTL set saliency, alternative hypothesis: “less”).

### Deletion screen

Separately, GM12878 DNase-seq and CTCF ChIP-seq peaks were merged if multiple peaks were found within 1500 bp of each other (bedtools merge -d 1500). Peaks were extended to at least 300 bp. All DNase peaks that overlapped with CTCF peaks were removed. For every remaining CTCF and DNase site we predicted the impact of deleting the respective site on chromatin interactions using the 5 kb GM12878 model. Chromatin classes were assigned based on overlap with the GenoSTAN chromatin segmentation described above.

### Additional software and packages

All neural networks were implemented in python (v3.5) and tensorflow (developed under 1.8.0; compatible with current latest stable version 1.14.0).

#### Additional Tools

- bedtools^44^
- deeptools (v2.4.2)^45^
- MACS (v2.0.10)^46^
- samtools (v1.3)^47^

Additional R packages
- cowplot (v0.6.2, https://github.com/wilkelab/cowplot)
- GenomicRanges – (v1.30.3)^48^
- GenoSTAN (STAN v2.6.0)^49^
- ggplot2 (v3.1.0)^50^
- RColorBrewer (v1.1.1-2, https://cran.r-project.org/web/packages/RColorBrewer/index.html)
- rtracklayer (v1.30.4)^51^
- tidyverse (https://www.tidyverse.org)
- zoo (v1.8.1)^52^

#### Additional Python libraries

- numpy (1.16.4)^52^
- h5py (v2.9.0, http://www.h5py.org)
- pysam (0.15.2, https://github.com/pysam-developers/pysam)

## Software availability

All code for training and employing deepC networks as well as trained models are available under: https://github.com/rschwess/deepC; All code for training and employing chromatin feature networks is available under: https://github.com/rschwess/deepHaem

## Acknowledgements

This work was supported by the MRC (reference MC_UU_12009/4) and the Wellcome Trust via Strategic Award (reference 106130/Z/14/Z) and Institutional Strategic Support Fund (reference 105605/Z/14/Z). R.S and R.B. are supported by the Wellcome Trust Genomic Medicine and Statistics PhD Programme (references 203728/Z/16/Z Ħ 203141/Z/16/Z). G.L. is supported by the Wellcome Trust supporting award (reference 090532/Z/09/Z). Y.W.T. is supported by the European Research Council under the European Union’s Seventh Framework Programme (reference FP7/2007-2013) ERC grant agreement no. 617071.

The Genotype-Tissue Expression (GTEx) Project was supported by the Common Fund of the Office of the Director of the National Institutes of Health, and by NCI, NHGRI, NHLBI, NIDA, NIMH, and NINDS. The data used for the analyses described in this manuscript were obtained from: the GTEx Portal on 01/03/19, release v7.

## Supplementary Figures

**Supplementary Figure 1.**
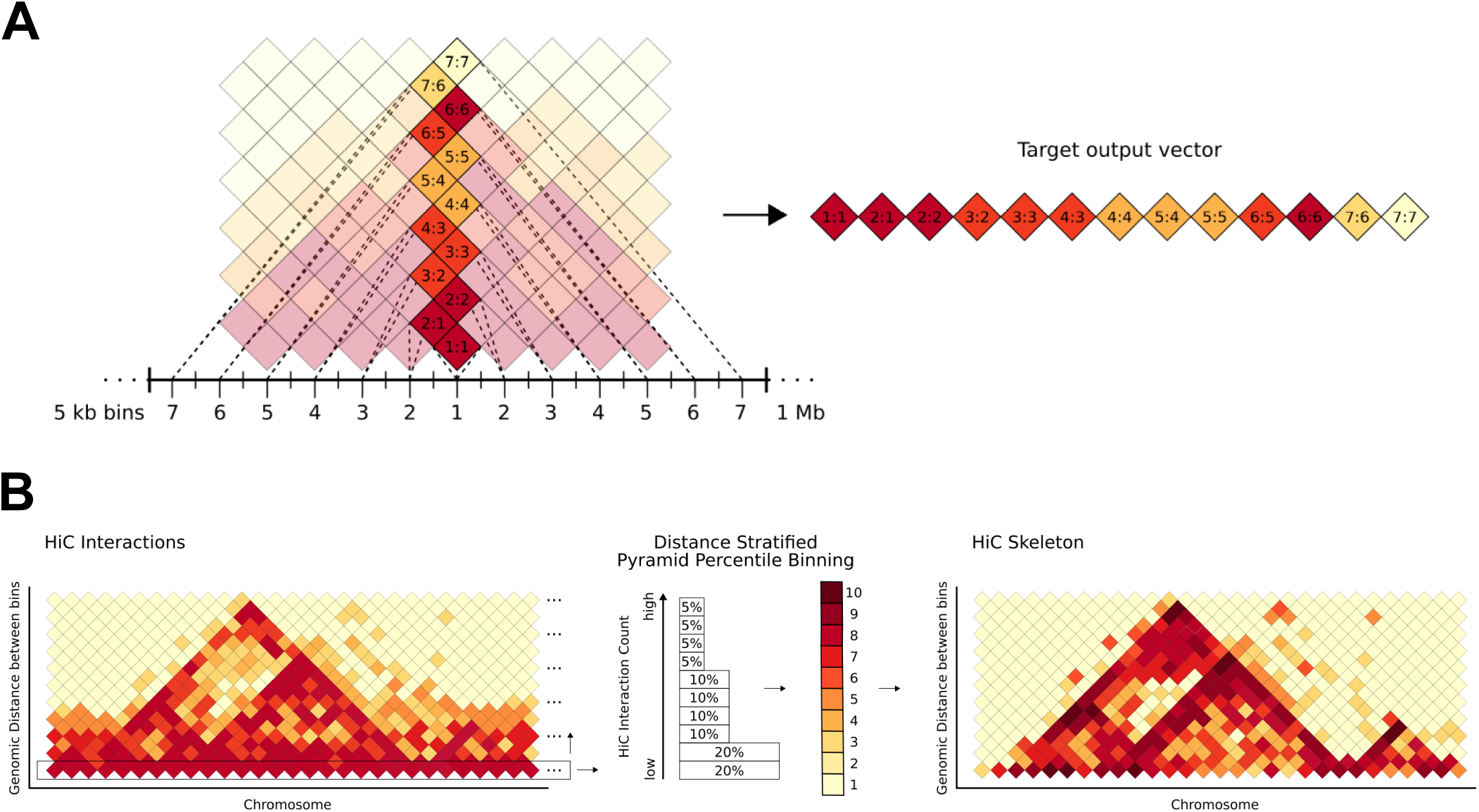
Scheme for encoding Hi-C data for deep learning. **A)** For a given window of length 1 Mb + 1x the bin size, the binned Hi-C data is recorded on a vertical (zig zag) pole centered in the center on the window, where each square represents the pairwise interaction strength of the bins. This is extracted as the target output vector for the given window. By moving the window over a chromosome this records all Hi-C interactions up to a genomic distance of 1 Mb between the bins. **B)** The Hi-C interactions are percentile binned in a genomic distance stratified manner. For every genomic distance, in steps equal to the bin size, the Hi-C signal is split into unequal percentiles ranging from 20 % bottom to 5 % top. The percentiles are attributed the values 1 to 10 yielding the Hi-C skeleton. The unequal percentile sizes ensure a finer distinction of the differences at the high Hi-C interaction value range, while minor differences in the low interaction value range are squished. Effectively, this procedure reduces the linear distance driven proximity signal and enhances domains and domains boundaries.

**Supplementary Figure 2.**
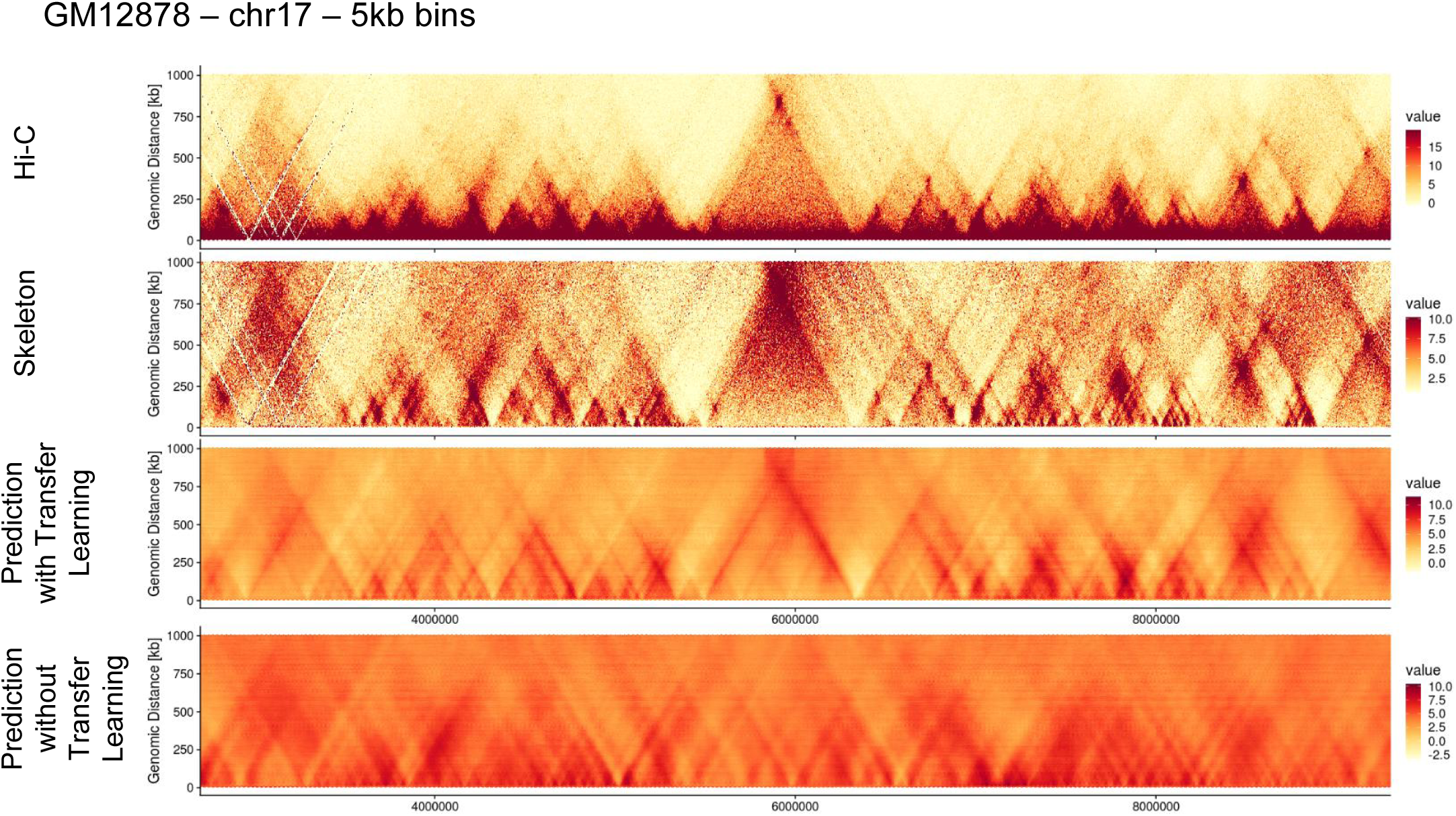
Comparison of deepC training with and without transfer learning. Training a deepC model with the same architecture but without pre-seeding the lower convolutional layers with the deepHaem model results in the emergence of triangular structures. Their positioning however is does not match with the Hi-C structures. In contrast, with pre-seeding the predicted domains overlap well with the Hi-C skeleton.

**Supplementary Figure 3.**
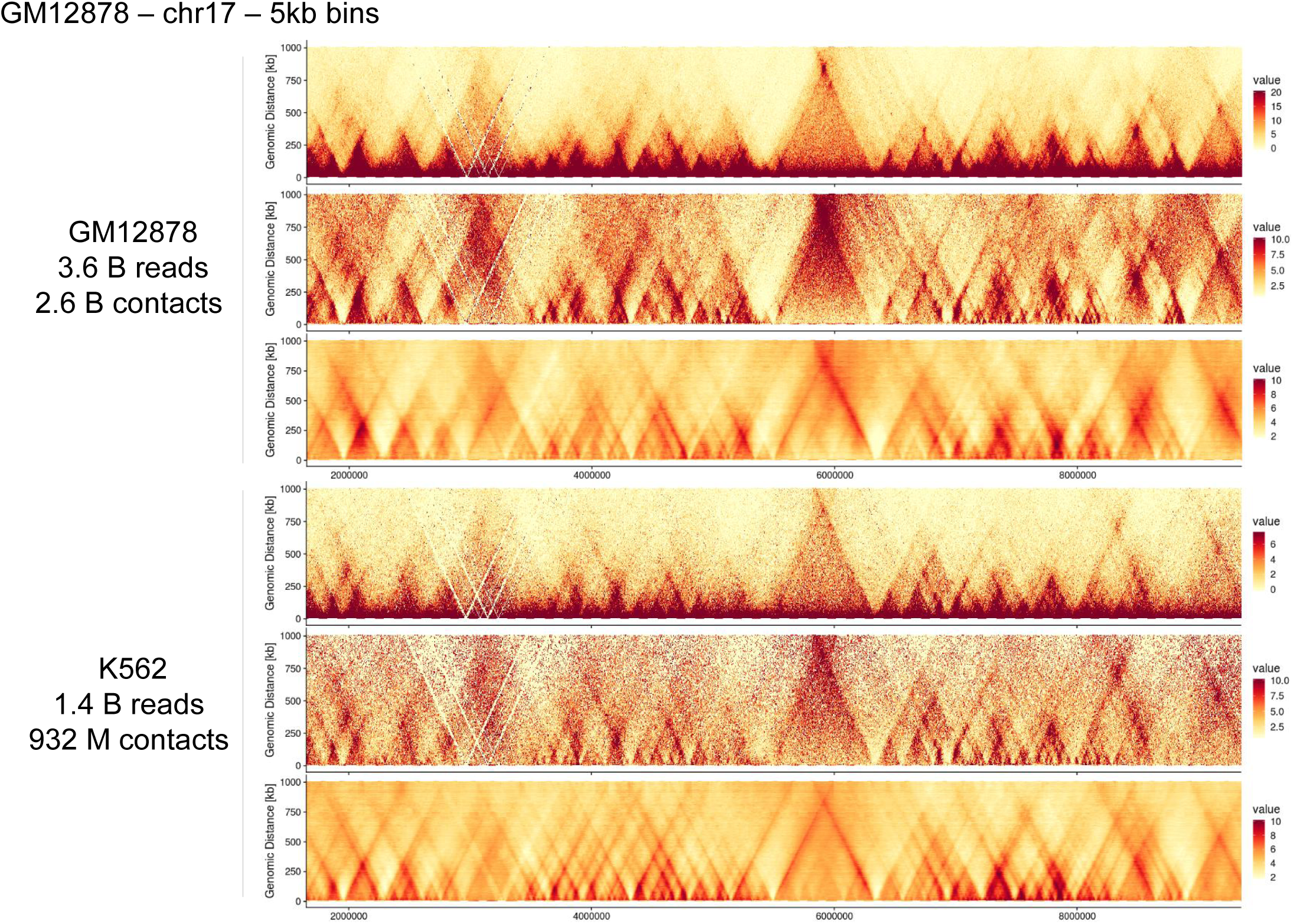
DeepC GM12878 and K562 example. Shown is a ∼ 7 Mb region on chr17 one of the test chromosomes.

**Supplementary Figure 4.**
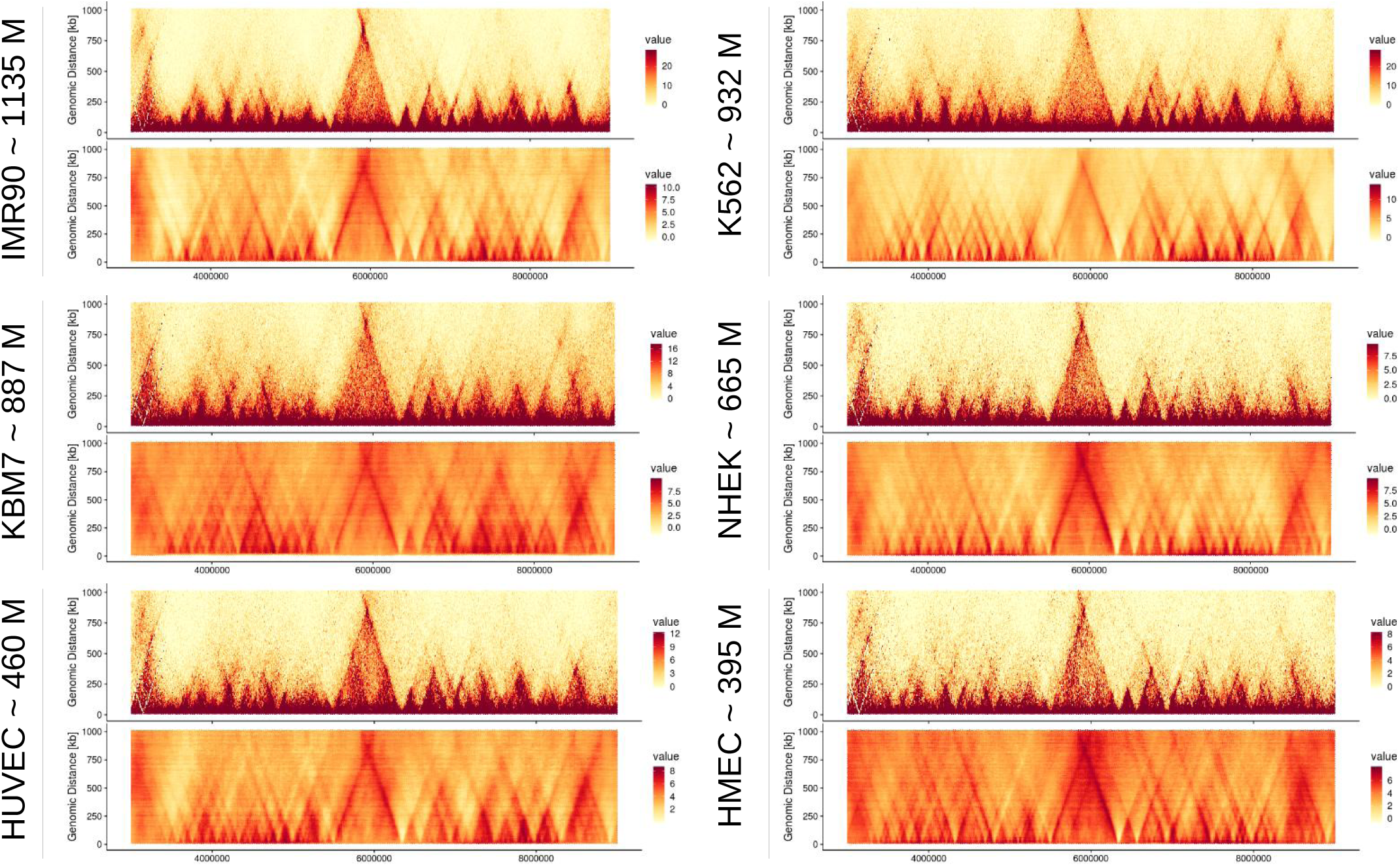
DeepC across 6 tissues. Shown are the Hi-C data and deepC prediction overlay. The number of valid Hi-C contacts per data set is recorded. Shown is a region ∼ 7 Mb on the validation chromosome chr17.

**Supplementary Figure 5.**
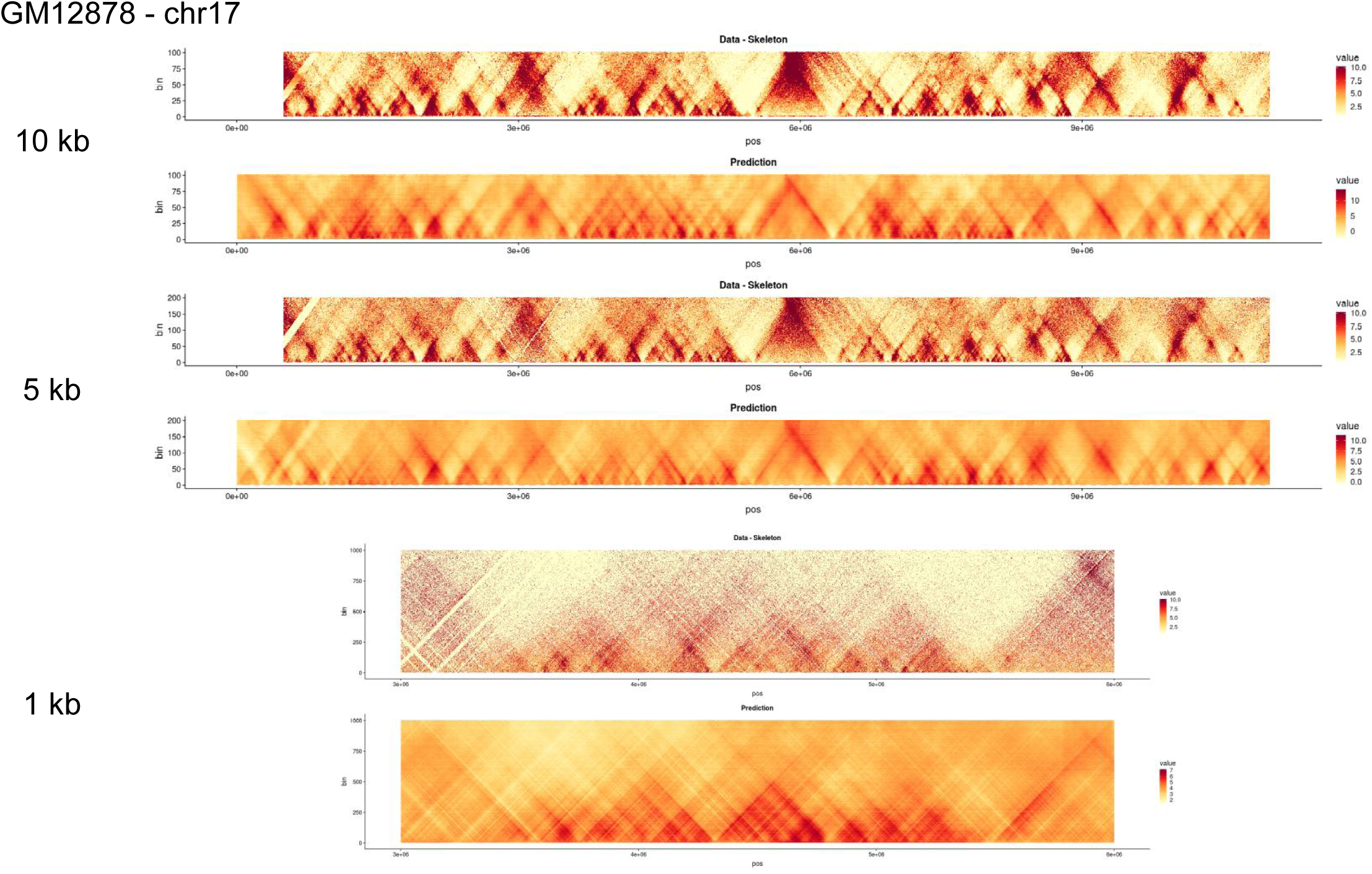
Training deepC at different bin sizes. Shown are pairs of Hi-C skeleton and deepC prediction at different bin size resolutions, all on validation chromosome 17.

**Supplementary Figure 6.**
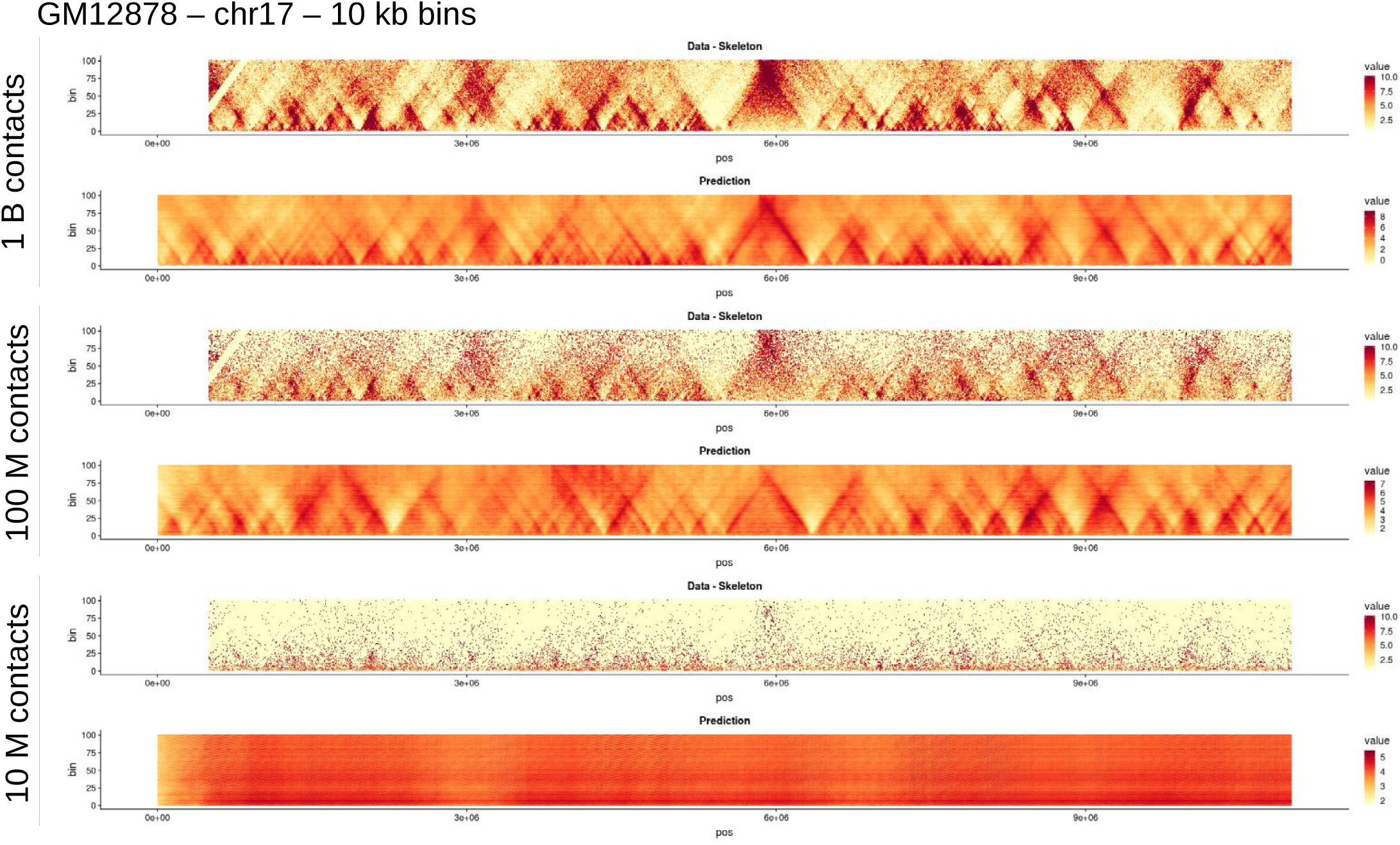
Effect of sub sampling HiC contacts on deepC predictions. We sub sampled the valid contacts of the realigned GM12878 HiC data (primary replicate with ∼36 B reads yielding ∼ 26 B valid contacts) to 1 B, 100 M and 10 M valid contacts, respectively. We then trained a deepC net at 10 kb resolution. For comparison, the cell type with the least amount of sequencing in Rao et al., HMEC is sequenced to ∼538 M reads yielding ∼395 M valid contacts.

**Supplementary Figure 7.**
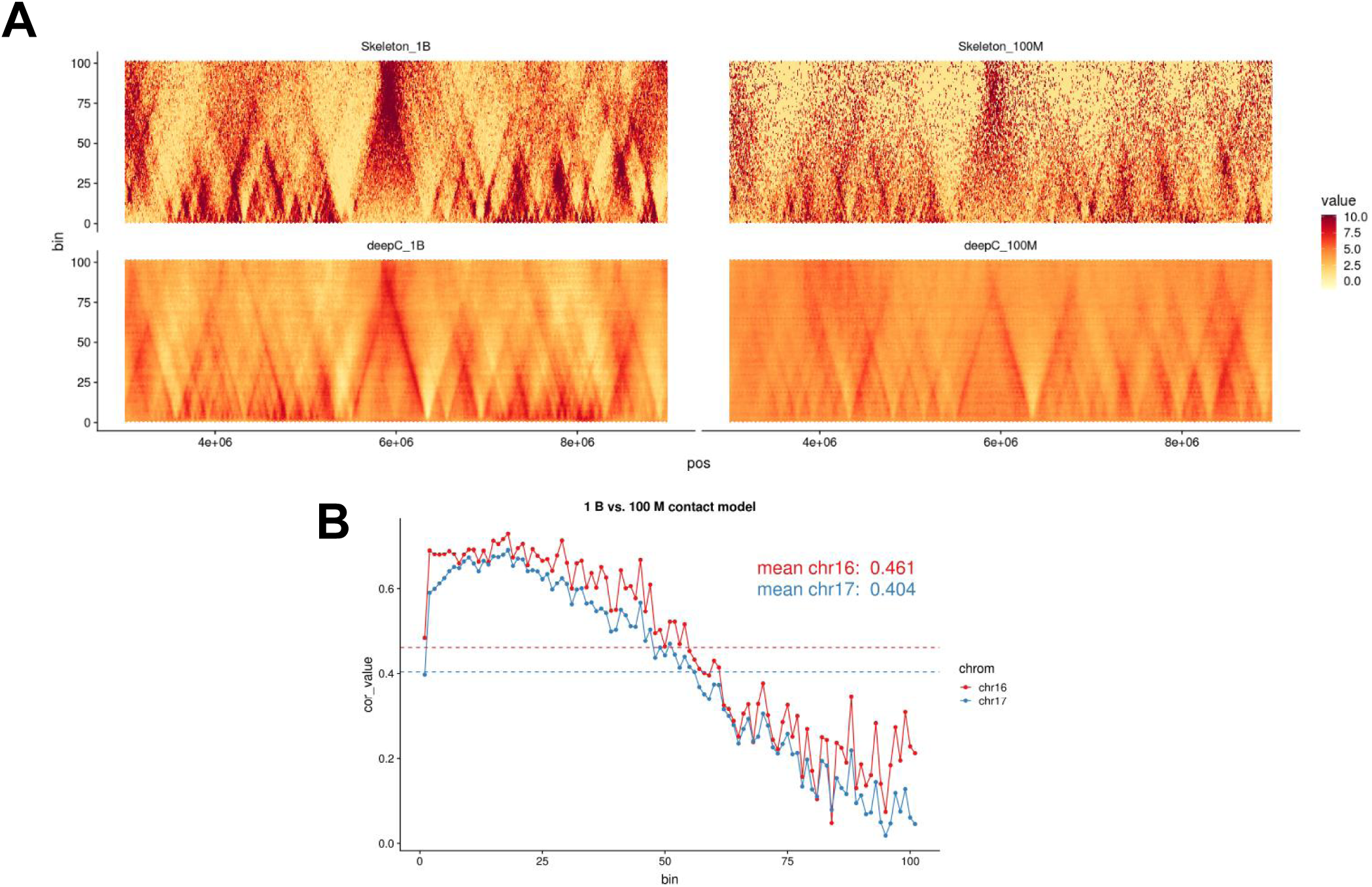
Effect of sub sampling HiC contacts on deepC predictions, detail with correlation. Distance bins are in multiples of the respective resolution, here 5kb. **A)** Detail of data shown in Supplementary Figure 6 above. **B)** The average Pearson correlation between the 1 B contact and the 100 M contact derived deepC prediction on validation chromosomes chr16 and chr17 was distance stratified. The correlation is stronger for shorter distances (∼ < 500 kb corresponding to distance bin 50) and drops for long ranged interactions. This suggests that deep sequencing better resolves long ranger interactions enabling deepC to learn to predict them more accurately. In contrast, shorter ranged interactions and smaller domains are resolved sufficiently for training and prediction.

**Supplementary Figure 8.**
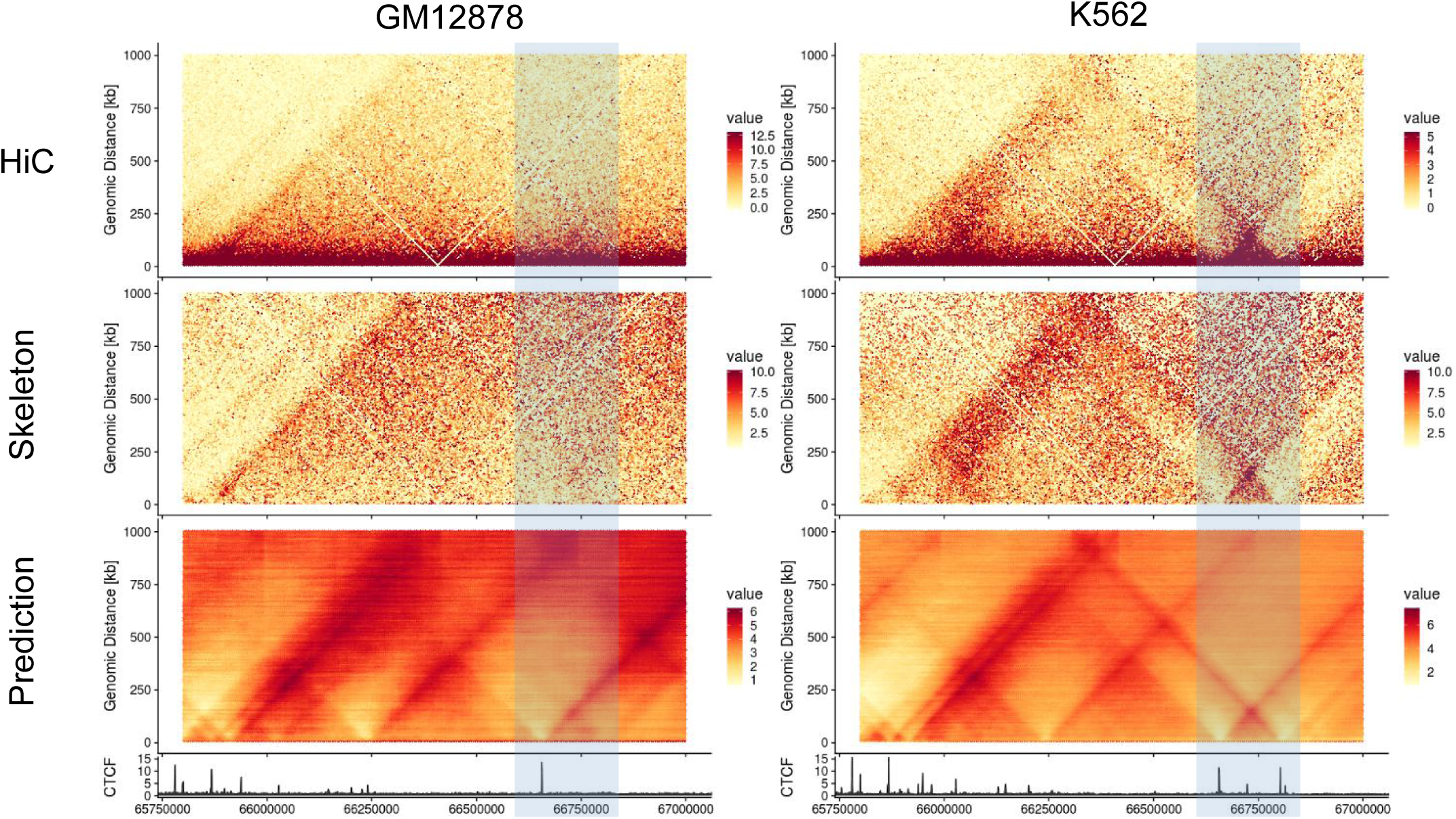
Tissue Specific deepC predictions. DeepC predicts sub TAD structure matching the tissue specific CTCF signals in K562. The subTAD is present in K562 HiC data.

**Supplementary Figure 9.**
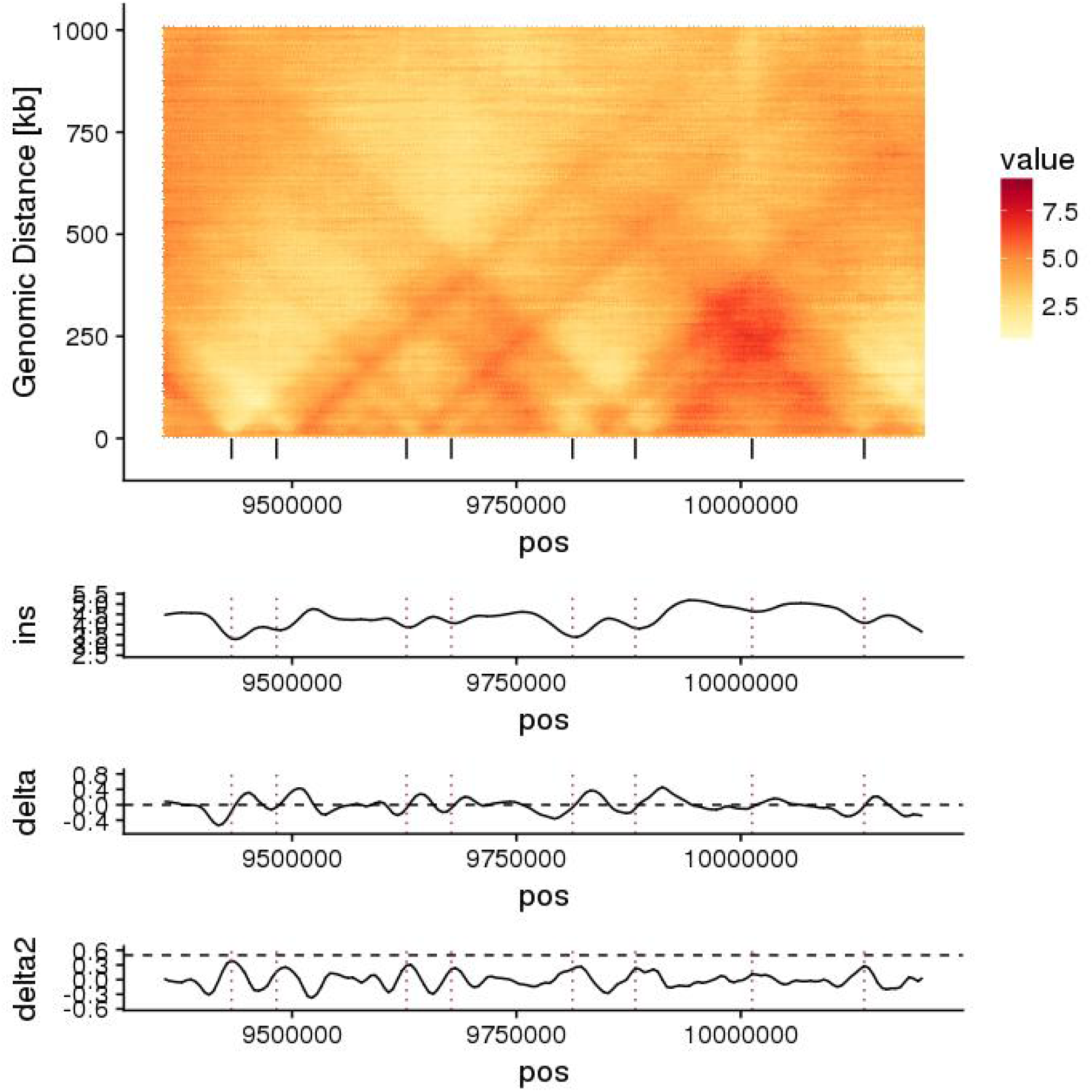
Insulation score based boundary calling. To map domains at high resolution we adapted the insulation score based calling method from Crane et al. 2015. We calculate the moving, mean insulation score (ins) using a window of 25 kb. We then calculate the approximation of the first and second order derivative (delta) and (delta2). Zero crossings in delta were identified and filtered on the a delta2 strength thresholds reflecting maxima in (ins) and the strength of boundaries as the change of change in insulation score (the sharpness).

**Suppl. Figure 10.**
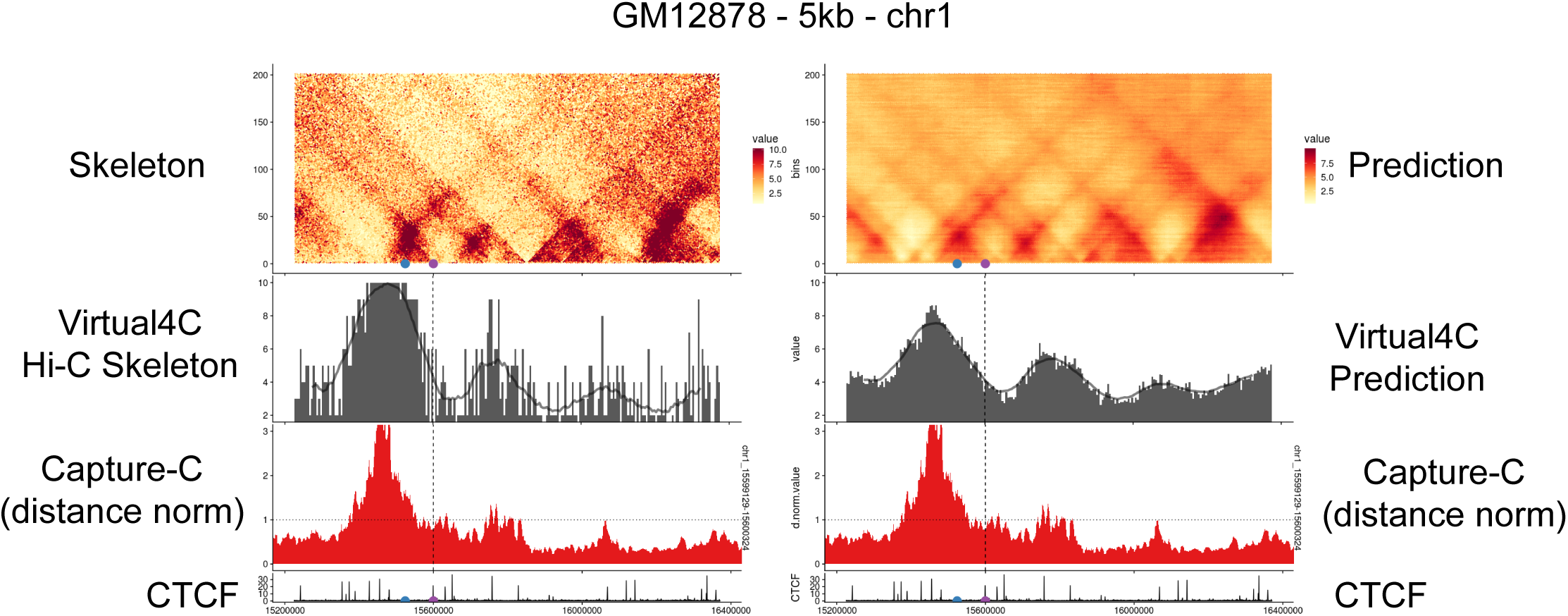
Virtual 4C vs. Capture-C. Shown are the Hi-C skeleton and the deepC prediction. From a viewpoint of interest, here a CTCF site (purple dot), we can derive a virtual 4C profile (grey) from the skeleton or the prediction respectively. The black line indicates a smoothed profile. The blue dot represents an intra domain viewpoint from which we captured as well (data not shown). We compare the virtual 4C profiles against the distance normalized Capture-C track (red) by calculating the Pearson correlation coefficient between the profiles.

**Suppl. Figure 11.**
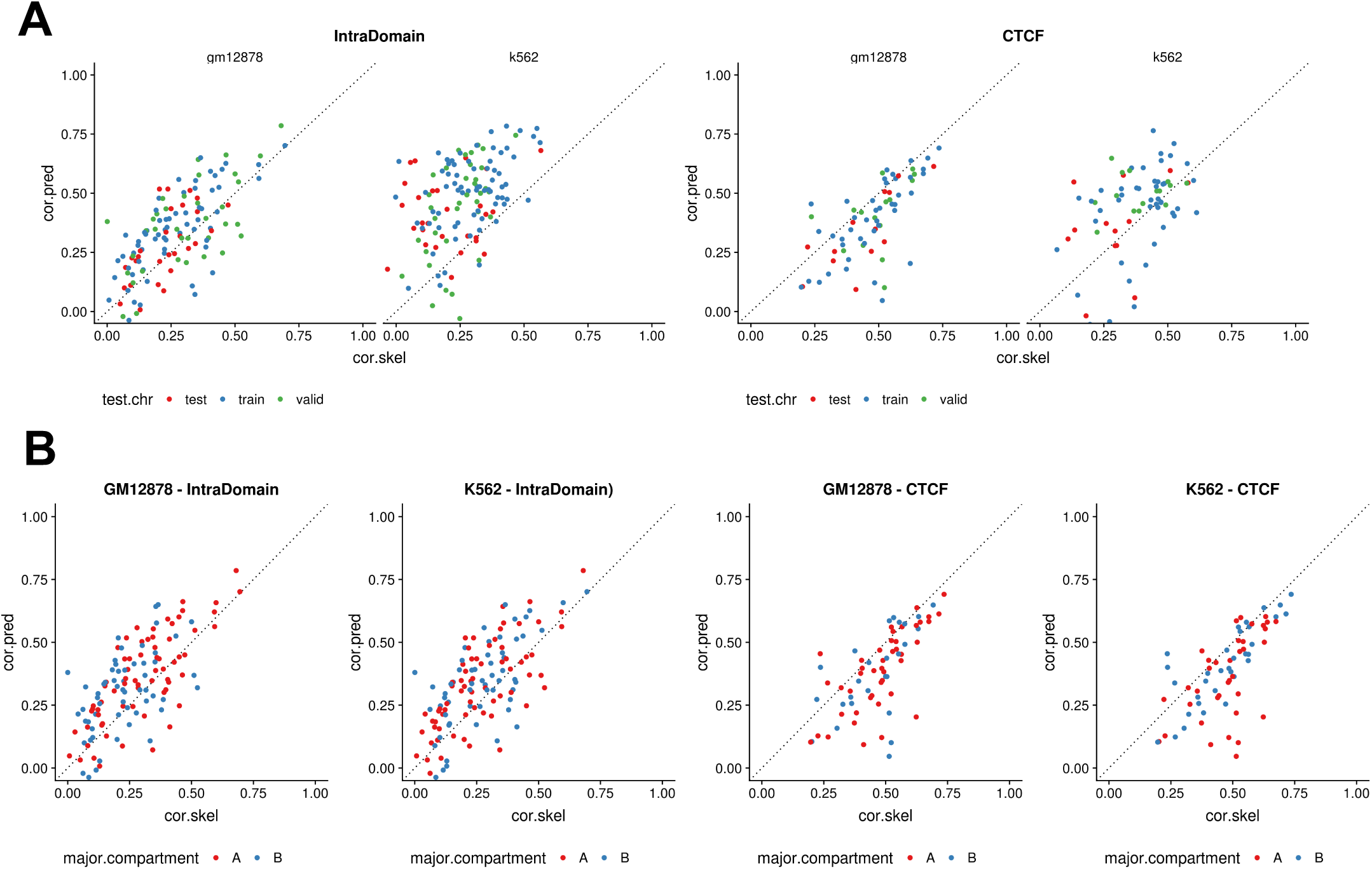
Stratifying virtual4C correlations. Shown is the Pearson correlation of the distance normalized Capture-C data with the virtual4C derived from the Hi-C skeleton and the deepC prediction map respectively for 220 viewpoints. The viewpoints have been stratified over **A)** location on train, test and validation set chromosomes and **B)** overlap with A and B compartments. No significant bias from these aspects can be observed.

**Supplementary Figure 12.**
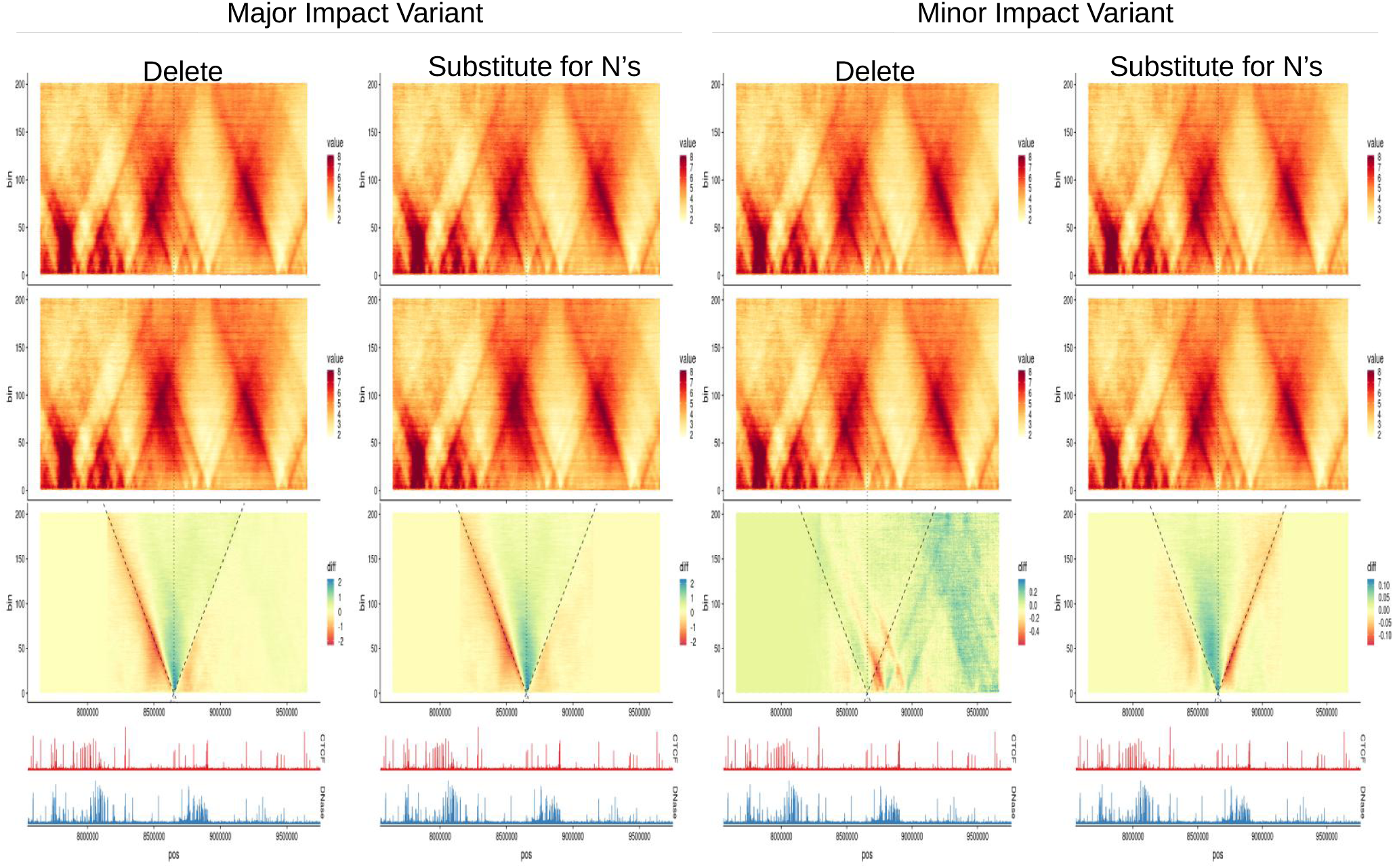
Effect of *in silico* deleting vs introducing N’s. Example of two *in silico* deletions spanning 100 bps each. Compared are the impact of deleting the respective sequence or replacing the sequence with a matching number of N’s. In major impact variants the shadow created by the shifting of the sequence downstream of the deletion is negligible. In low impact variants the shadow becomes the dominating feature.

