## Supplementary Notes for "DeepC: Predicting chromatin interactions using megabase scaled deep neural networks and transfer learning"

### Supplementary Note

##### Chromatin Feature Network Architecture and Hyperparameters

1) Five convolutional (conv.) layers with ReLU activation followed 1D max pooling and dropout in the following scheme

| Layer | Hidden Units | Filter Width | Max Pool Width |
|---|---|---|---|
| 1 | 300 | 8 | 4 |
| 2 | 600 | 8 | 5 |
| 3 | 600 | 8 | 5 |
| 4 | 900 | 4 | 5 |
| 5 | 900 | 4 | 2 |

2) Fully connected layer with sigmoid activation

Additional training parameters:

- Batch size: 100
- Dropout Probability: 0.2
- Learning Rate (Intital): 0.0001
- L2 regularizer strength: 0.001
- Initializer: Xavier (Glorot & Bengio, 2010) for filters (tf.constant_initializer for biases)
- (ADAM) Beta 1: 0.9
- (ADAM) Beta 2: 0.999
- (ADAM) Epsilon: 0.1
- Shuffle training set after every epoch
- Total trainable parameters: 10572000

##### Chromatin Interaction Network Architecture and Hyperparameters

1) Five conv. layers with ReLU activation followed 1D max pooling and dropout (exact architecture as conv. layers in the chromatin feature network). Filter weights and biases are seeded with the weights from the trained chromatin feature network. Same dropout applied.

2) The conv. layers are followed by a 1x1 conv. layer that maps the 900 hidden units of the last conv. layer to the 100 dilational conv. units for the following layers.

2) 9 Dilated conv. layers. 100 hidden units per layer and a dilation width of 3. The scheme of dilation rates was designed to reach the full 1 Mb + 1 x HiC bin size used (e.g. 1005000 for 5 kb network), doubling the dilation rate every layer. Dilation scheme: 2,4,8,16,32,64,128,256,1. The last layer with a dilation rate 1 was introduced to ensure equal coverage of every position along the sequence axis at the bottom layer (see Supplementary Note Figure 1). Following PixelCNN (Oord, Kalchbrenner, & Kavukcuoglu, 2016) and WaveNet (Oord, Dieleman, et al., 2016), and its implementation in (https://github.com/ibab/tensorflow-wavenet), we chose to use gated conv. layers. Gated conv. layers require training double the amount of filter parameters, but have the potential of modelling more complex functions through the multiplicative units, similar to LSTMs. For every dilated unit in every layer, a separate filter and gate with equal dimensions are trained. A hyperbolic tangent is used as activation for the filter and a sigmoid activation for the gate. Filter and gate outputs are then multiplied element wise. (see Supplementary Note Figure 2). We also adapted residual connections between the dilated layers.

3) Fully connected layer. The output of the fully connected layer are the predicted interaction strengths (in units matching the pyramid percentile normalization). The model is trained using the ADAM optimizer, to minimize the mean square error between the outputs and the true labels.

Additional training parameters:

- Batch size: 1
- Dropout Probability: 0.2
- Learning Rate (Intital): 0.0001
- L2 regularizer strength: 0.001
- Initializer: Seedeing + Xavier (Xavier Glorot and Yoshua Bengio (2010)) for filters (tf.constant_initializer for biases)
- (ADAM) Beta 1: 0.9
- (ADAM) Beta 2: 0.999
- (ADAM) Epsilon: 0.1
- Shuffle training set after every epoch

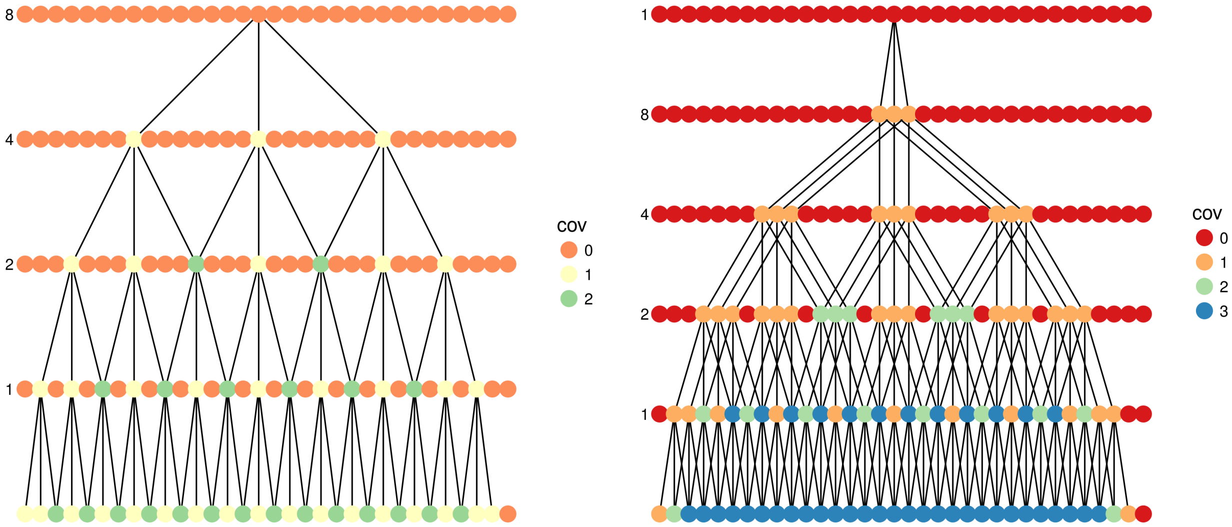


**Supplementary Note Figure 1.** Illustration of the effect of a final, additional dilation layer with dilation rate 1 on the coverage per element along the sequence axis. Lines of points represent a hidden dilated layer with every point being an element along the sequence axis. The dilation rate per layer are indicate adjacent. The colors indicate the coverage, the amount of of connections to downstream units at a single time step.

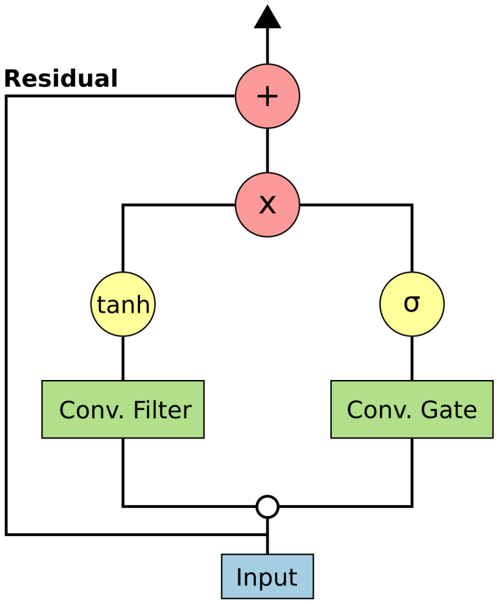


**Supplementary Note Figure 2.** Scheme of a single dilated, gated conv. unit with residual.

#### Supplementary Tables

**Supplementary Table 1. Datasets used for training the chromatin feature network.**
Please see separate table file.

**Supplementary Table 2. Capture-C validation probes.**
Please see separate table file.

**Supplementary Table 3. Distance thresholds in bp.** Interactions below 2500 bps were excluded. Everything below the near threshold was classed as near and so on. Everything above the far thresholds was excluded from the distance fit.

| Set | Near [bp] | Intermediate [bp] |
|---|---|---|
| GM12878 CTCF | 125000 | 125000 |
| GM12878 Intra Domain | 125000 | 1250000 |
| K562 CTCF | 170000 | 3250000 |
| K562 Intra Domain | 40000 | 3500000 |

**Supplementary Table 4. Auxiliary ENCODE chromatin data sets used for chromatin segmentation and visualization.**

| Experiment ID | Cell Type | Assay |
|---|---|---|
| ENCSR000EMT | GM12878 | DNase-seq |
| ENCSR000AKB | GM12878 | CTCF ChIP-seq |
| ENCSR057BWO | GM12878 | H3K4me3 ChIP-seq |
| ENCSR000AKF | GM12878 | H3K4me1 ChIP-seq |
| ENCSR000AKC | GM12878 | H3K27ac ChIP-seq |
| ENCSR921NMD | K562 | DNase-seq |
| ENCSR000AKO | K562 | CTCF ChIP-seq |
| ENCSR668LDD | K562 | H3K4me3 ChIP-seq |
| ENCSR000AKS | K562 | H3K4me1 ChIP-seq |
| ENCSR000AKP | K562 | H3K27ac ChIP-seq |
